# Developmental Control of Long-Distance mRNA Transport Involves m^5^C RNA Methylation and ALYREF Nuclear Export Factors

**DOI:** 10.1101/2024.05.30.596576

**Authors:** Ying Xu, András Székely, Steffen Ostendorp, Saurabh Gupta, Lei Yang, Federico Apelt, Yağmur Hasbioğlu, Yan Zhao, Eleni Mavrothalassiti, Linda Wansing, Eleftheria Saplaoura, Friedrich Kragler

## Abstract

In grafted plants mRNAs can be transported from shoot to root, however, how their mobility is regulated remains poorly understood. Recent work has shown that m^5^C methylation plays an essential role in systemic mRNA mobility in vegetative plants, but its role at later developmental stages remains unclear. To address this, we examined the mobility of three mobile mRNAs, *TCTP1*, *HSC70.1* and *GRP7* in mutants deficient in m^5^C mRNA methylation. By comparing vegetative and flowering stages, we found that in contrast to *GRP7,* the reduced mobility of *TCTP1* and *HSC70.1* observed in the vegetative growth phase is restored upon flowering, indicating that m^5^C-dependent regulation of RNA transport is developmentally controlled. We further identified two RNA-binding proteins, the nuclear mRNA export factors ALY2 and ALY4, as phloem mobile. *ALY2* and *ALY4* deficient plants show a similar developmental effect on mRNA mobility as m^5^C mRNA methylation mutants, suggesting that they contribute to the regulation of systemic mRNA transport. Notably, combined perturbation of m^5^C methylation and ALY2 or ALY4 results in reduced mRNA mobility even in flowering plants. Together, our findings reveal an unexpected developmental layer of control over mRNA transport and identify the nuclear mRNA export factors ALY2 and ALY4 as potential regulators of long-distance RNA movement.

**SIGNIFICANCE STATEMENT:** mRNA m^5^C methylation is essential for shoot-to-root mRNA transport during plant growth; however, whether and how mRNA mobility is regulated, and whether it changes during development remains unclear. We show that *TCTP1* and *HSC70.1* mRNA transport is reduced in vegetatively growing m^5^C-deficient mutants but remains unaffected upon flowering, and identify ALY2 and ALY4 proteins, two nuclear export factors, as additional regulators of long-distance mRNA transport, providing new layer of mRNA transport.

## INTRODUCTION

The systemic transport of mRNAs across plant tissues is a fundamental mechanism underlying long-distance signaling, developmental coordination, and environmental adaptation. While the existence of graft-mobile transcripts has been well documented, the molecular mechanisms governing their selection, export, and delivery remain poorly understood. A wide variety of small RNAs and transcripts larger than 100 bases such as messenger RNAs (mRNA), viral RNAs (vRNA), and non-coding RNAs (ncRNA) are systemically transported to distant tissues in plants (Zhang *et al*., 2014, Li *et al*., 2021, Cheng *et al*., 2023). Local mRNA transfer between cells appear to depend on RNA-binding proteins as shown with RIBOSOMAL RNA-PROCESSING PROTEIN 44A (AtRRP44A), which interacts with intercellularly transported *KNOTTED1 and STM* homeodomain transcripts in *Arabidopsis thaliana* (*A. thaliana*) (Kitagawa *et al*., 2022). Graft-mobile transcripts such as *FLOWERING LOCUS T* (*FT*), *BEL5*, *TRANSLATIONALLY CONTROLLED TUMOR PROTEIN 1* (*TCTP1*), *CHOLINE KINASE 1* (*CK1*) and *HEAT SHOCK COGNATE PROTEIN 70.1* (*HSC70.1*) appear to serve as systemic signals, regulating plant development and growth (Banerjee *et al*., 2006, Lu *et al*., 2012, Thieme *et al*., 2015, Yang *et al*., 2015, Zhang *et al*., 2016, Ghate *et al*., 2017, Xia *et al*., 2018, Yang *et al*., 2019). For a small number of graft-mobile transcripts, structural features enabling their transport to distant tissues have been identified (Huang and Yu, 2009, Zhang *et al*., 2016, Yang *et al*., 2019, Yang *et al*., 2023a, Yang *et al*., 2023b). For example, relatively small RNA regions present in *GIBBERELLIC ACID INSENSITIVE (GAI)* (Huang and Yu, 2009), *TCTP1* (Yang *et al*., 2019), and *HSC70.1* (Yang *et al*., 2023b) are necessary for their transport from shoot to root. Furthermore, a co-transcribed tRNA-Gly sequence present at the 3’ UTR of *CK1* mRNA was shown to be necessary for shoot-to-root and root-to-shoot long-distance transport. Such tRNA-like sequences (TLS) are also mediating long-distance transport of heterologous fusion transcripts (Zhang *et al*., 2016, Yang *et al*., 2023a). In addition, secondary 5-methylcytosine (m^5^C) modifications seem to be necessary for *HSC70.1* and *TCTP1* transcript transport across graft junctions in vegetative *A. thaliana* (Yang *et al*., 2019) and for *CK1* in pumpkin (Li *et al*., 2024). The evidence to date thus points towards structural motifs and secondary mRNA modifications being important determinants of mRNA mobility (Li *et al*., 2024). However, how mRNAs are selected for delivery and whether mRNA transport is regulated differently in adult (committed to flowering) or juvenile (vegetative) plants is not well understood.

RNA methylation is a dynamic and reversible process. Methyltransferases (writers) target specific mRNAs, RNA-binding proteins (readers) recognize the modified transcripts, and demethylases (erasers) remove the modifications (Meyer and Jaffrey, 2014, Xue *et al*., 2020). RNA methylation occurs within mRNA (Cui *et al*., 2017, David *et al*., 2017), tRNA, ribosomal RNA (rRNA), and long non-coding RNA in mammals and plants (Goll *et al*., 2006, Burgess *et al*., 2015, Zhang *et al*., 2016, David *et al*., 2017, Gao and Fang, 2021). In plants, m^6^A is the most abundant mRNA modification, playing essential roles in various developmental processes such as floral transition, trichome branching and shoot growth, and also in response to biotic and abiotic stress, by affecting mRNA processing and metabolism (Shen *et al*., 2016, Duan *et al*., 2017, Scutenaire *et al*., 2018, Wei *et al*., 2018, Zhang *et al*., 2019, Song *et al*., 2021, Zhou *et al*., 2021, Hou *et al*., 2022, Amara *et al*., 2023, Prall *et al*., 2023). Similarly, m^5^C plays a role in root development and responses to abiotic stresses, such as heat and oxidative stress (Cui *et al*., 2017, David *et al*., 2017, Tang *et al*., 2020). The best studied RNA cytosine methyltransferases are NSUN2B, a member of the human NOP2/Sun (NSUN) domain family also known as tRNA METHYLTRANSFERASE 4B (TRM4B; TAIR# AT2G22400), and DNMT2, also known as tRNA ASPARTIC ACID METHYLTRANSFERASE 1 (TRDMT1; TAIR# AT5G25480) (Goll *et al*., 2006, Yang *et al*., 2017), which have been validated as m^5^C writers (Apelt *et al*., 2017, Cui *et al*., 2017, Yang *et al*., 2019, Tang *et al*., 2020). Arabidopsis NSUN2B (TRM4B) mutants show reduced mRNA and tRNA methylation whereas the NSUN2B paralog TRM4A appears be catalytically inactive (Burgess *et al*., 2015, David *et al*., 2017). Although three m^5^C readers, namely the highly conserved RNA-binding proteins ALYREF, Y-box binding protein 1 (YBX1) and RAD52, have been identified in mammalian cells (Yang *et al*., 2017, Chen *et al*., 2019, Yang *et al*., 2019), plant-related m^5^C readers have yet to be confirmed in their *in planta* function.

Exon junction complex proteins, such as ALYREFs are part of the nuclear transcript export complex (TREX) and link mRNA transcription, splicing with nuclear export via the nuclear pore complex. The *A. thaliana* ALYREF family has four members: ALY1, ALY2, ALY3, and ALY4. No obvious phenotypes were reported for *aly* single and double mutants, whereas *aly* quadruple mutants were delayed in bolting and reduced in seed production (Pfaff *et al*., 2018a). All four ALY family members were shown to be involved in mRNA nuclear export by interacting with the DEAD-box RNA helicase UAP56 (Pfaff *et al*., 2018a). Although all four ALYs interact with the siRNA transport-limiting P19 protein produced by Tomato Bushy Stunt Virus, only ALY2 and ALY4 exhibited cytoplasmic retention in the presence of the P19 siRNA-binding protein (Uhrig *et al*., 2004), implying that cytosolic ALYs may contribute to the systemic delivery of antiviral siRNAs and/or of viral RNAs. ALY1 was shown to preferentially bind to m^5^C oligonucleotides suggesting that ALY family members are potential m^5^C readers (Pfaff *et al*., 2018a). Notably, ALY2 and ALY4 proteins interact with each other, ALY2 also binds to the microtubule organizing CONVOLUTA/SPIRAL 2 and HAPLESS 1/MAGO proteins, and ALY4 also binds to the exon junction complex core protein eLF4A-III (Koroleva *et al*., 2009, Arabidopsis Interactome Mapping, 2011).

Here, we show that the reported loss of *HSC70.1* and *TCTP1* mRNA shoot-to-root transport in vegetative *dnmt2 nsun2b* m^5^C methyltransferase mutant plants (Yang *et al*., 2019) does not hold in flowering plants after flower initiation, i.e. transport is reinstated despite deficiency in m^5^C methylation. To gain further insights into the regulatory factors deciding on mRNA delivery from shoot to root, we analyzed the potential role of the two putative m^5^C readers, namely ALY2 and ALY4, and of DNMT2 and NSUN2B methyltransferases in mRNA transport in flowering committed versus vegetative plants. mRNA transport assays with *aly2* and *aly4* single mutants, as well as *aly2 dnmt2 nsun2b* and *aly4 dnmt2 nsun2b* triple mutants suggest that the two nuclear RNA export factors act together with DNMT2 and NSUN2B methyltransferases to facilitate mRNA delivery to roots. We propose that ALY2 and ALY4 protein facilitate cellular export of mRNAs via delivering intercellular mobile mRNAs to and via plasmodesmata suggested to contain nuclear pore proteins (Schladt *et al*., 2025) to enter the phloem.

## RESULTS

### *HSC70.1* and *TCTP1* Mobility is Restored in *dnmt2 nsun2b* Mutants Committed to Flowering

In vegetative *A. thaliana* plants, shoot-to-root transport of *TCTP1* and *HSC70.1* mRNA depends on m^5^C methylation. This was previously demonstrated using vegetative *dnmt2 nsun2b* mutants, which exhibit reduced m^5^C methyltransferase activity (Yang *et al*., 2019). To further examine this relationship across developmental stages, we compared the mobility of *TCTP1* and *HSC70.1* transcripts in grafted vegetative and flowering plants (Figure 1). Transgenic *35S::YFP-TCTP1* and *35S::YFP-HSC70.1* lines in Col-0 (wild-type) or *dnmt2 nsun2b* backgrounds were used as scions and grafted onto Col-0 or *dnmt2 nsun2b* rootstocks. Transcript mobility was assessed by RT-PCR (Figure 1B and C). Consistent with previous reports (Yang *et al*., 2019, Yang *et al*., 2023b), *YFP-TCTP1* and *YFP-HSC70.1* transcripts produced in *dnmt2 nsun2b* scions were not detected in the roots of vegetative plants (Figure 1B and 1C). In contrast, when these grafted plants developed to the flowering stage, shoot-to-root transport of both transcripts was restored (Figure 1B and 1C).

**Figure 1:**
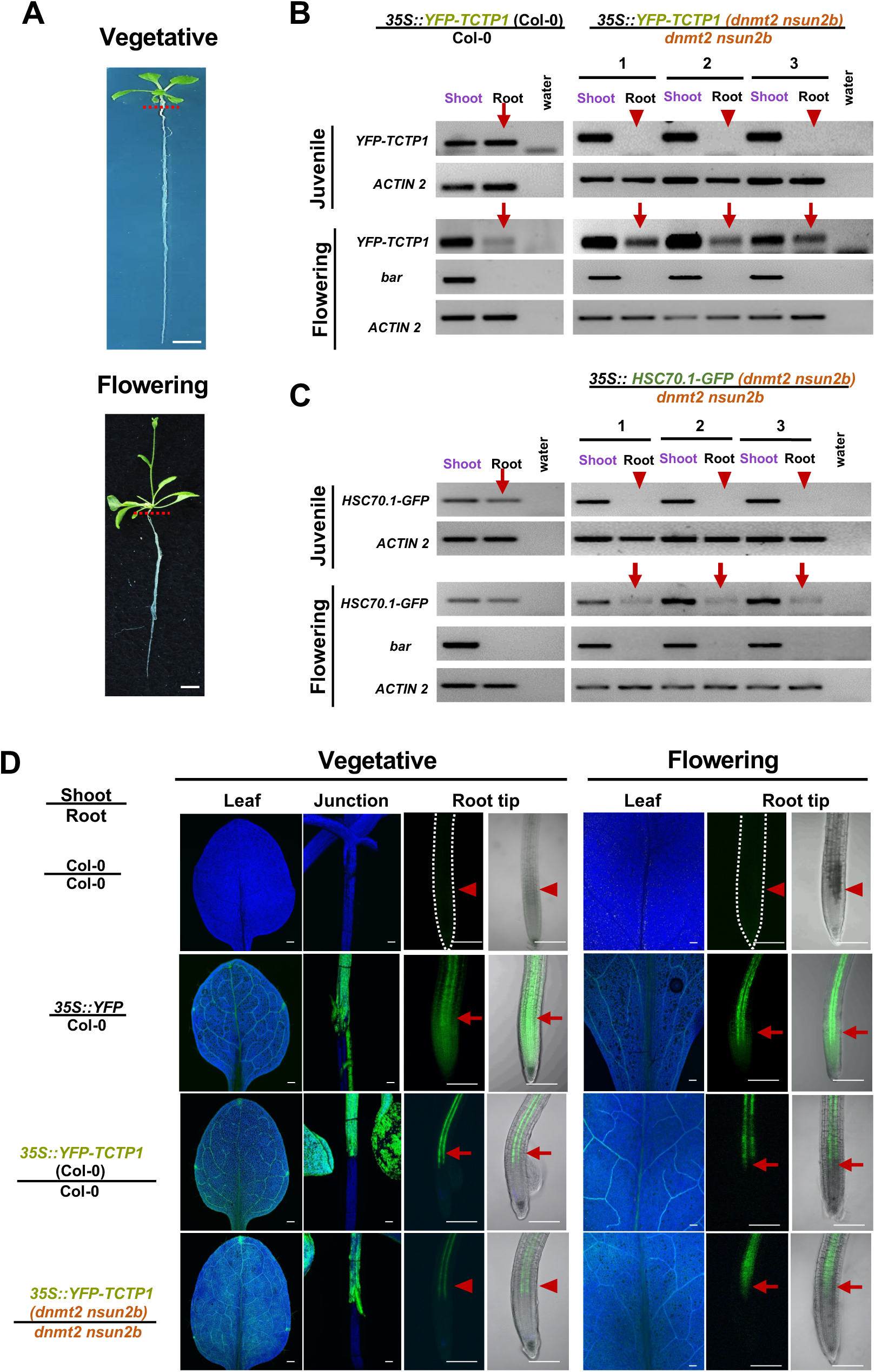
mRNA mobility in grafted vegetative vs. flowering Arabidopsis. (A) Representative images of grafted vegetative or flowering plants used to test mRNA transport. Scale bar: 1 cm. (B and C) RT-PCR assays to detect *YFP-TCTP1* (B) and *HSC70.1* (C) in shoot and root RNA samples of grafted vegetative and flowering plants. Arrowheads indicate absence, arrows indicate presence of *YFP-TCTP1*, *HSC70.1-GFP* and *GRP7* in root samples of grafted plants. For *YFP-TCTP1*, *HSC70.1-GFP* and *GRP7* detection in roots, 45 PCR cycles were used to detect mobile mRNA. *bar* (BastaR) and *nptII* specific primers were used for sample contamination controls (45 cycles) and *ACTIN2* specific primers were used for cDNA quality control (30 cycles). For RNA sample preparation, scions and roots of grafted vegetative (n = 10) or flowering (n = 3) plants was pooled. (D) YFP fluorescent signal detected by confocal laser scanning microscope (CLSM) in leaves, graft junctions, primary root tips of grafted plants. Note the lower YFP-TCTP1 signal detected in roots of grafted vegetative *dnmt2 nsun2b* mutant plants compared to wild-type and mutant flowering plants. Blue: auto-fluorescent background, arrows: YFP presence, arrowheads indicate weak or no YFP signals in roots of grafted plants. Scale bars: 200 µm.

To exclude nonspecific RNA movement, we examined the mobility of heterologous *YFP* mRNA expressed from the *35S* promoter. Consistent with previous report (Haywood *et al*., 2005, Paultre *et al*., 2016, Yang *et al*., 2019), *YFP* transcripts were not detected in roots of either vegetative or flowering plants (Figure S1A). In addition, endogenous *HSC70.1* transcripts are graft-mobile (Yang *et al*., 2023b), indicating that mobility of the *YFP* fusion transcripts depends on the *TCTP1* or *HSC70.1* sequence rather than the *YFP* tag. In line with transcript mobility, YFP-TCTP1 protein accumulation in grafted roots was strongly reduced in vegetative *dnmt2 nsun2b* plants but restored in flowering plants (Figure 1D). In wild-type vegetative roots, YFP-TCTP1 fluorescence was primarily detected in phloem pole pericycle (PPP) cells, with weaker signals in adjacent endodermal cells and no signal in xylem pole pericycle (XPP) cells (Yang *et al*., 2019). A similar distribution was observed in flowering plants (Figure S1B), indicating that the recipient cell types remain unchanged across developmental stages.

Overall, these results show that shoot-to-root transport of *HSC70.1* and *TCTP1* mRNAs depends on DNMT2- and NSUN2B-mediated m^5^C methylation in vegetative plants but not in flowering plants.

### m^5^C methyltransferase-mediated mRNA transport is floral-transition dependent

To determine whether the restored mRNA mobility in flowering *dnmt2 nsun2b* plants is driven by plant age or by the floral transition, we used an inducible flowering system. We generated *dnmt2 nsun2b ft* triple mutants expressing *35S::YFP-TCTP1* and a β-estradiol–inducible *SUC2::FT-GFP* construct, allowing controlled activation of florigen expression and floral transition (Liu *et al*., 2019a). When floral transition was induced in vegetative plants by β-estradiol treatment, FT-GFP protein was detected in both shoots and roots, confirming activation of the flowering program (Figure 2A). However, *YFP*-*TCTP1* transcripts remained undetectable in roots (Figure 2B), indicating that initiation of flowering is not sufficient to restore *YFP*-*TCTP1* transport.

**Figure 2:**
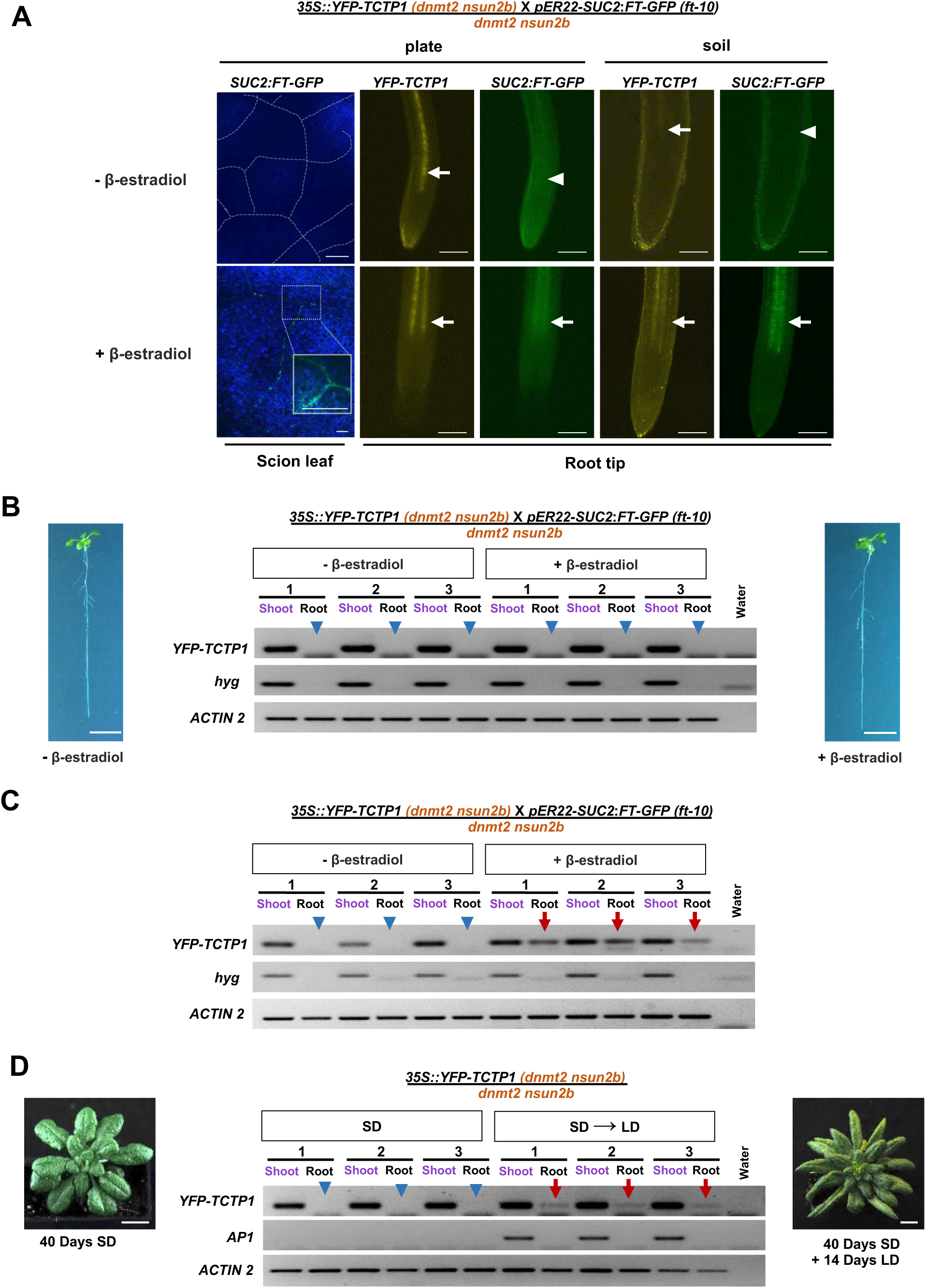
Mobility of *YFP-TCTP1* before and after flowering. (A) FT-GFP and YFP-TCTP1 fluorescence in leaves and roots of grafted plants with or without β-estradiol induction. For plate-grown plants, β-estradiol was applied 10 days after grafting and samples were analyzed after 2 days under short-day (SD) conditions (left). For soil-grown plants, grafted plants were transferred to soil 10 days after grafting. β-estradiol treatment was applied 7 days after transfer under long-day (LD) conditions and plants were analyzed two weeks later, after the onset of bolting (right). Blue, autofluorescence. Arrows indicate presence of FT-GFP or YFP-TCTP1; arrowheads indicate absence of FT-GFP signal in roots. Scale bars, 200 µm. (B and C) RT-PCR detection of *YFP-TCTP1* transcripts in shoots and roots of grafted plants from (A) grown on plate (B) or soil (C). (D) RT-PCR detection of *YFP-TCTP1* transcripts in shoots and roots of grafted plants grown under SD conditions for 40 days or shifted from SD (40 days) to LD conditions for 14 days. For *YFP- TCTP1* detection 45 PCR cycles were used. *hyg* transcript specific primers were used for contamination detection (45 cycles). *ACTIN2* specific primers were used for cDNA quality control (30 cycles). For floral meristem appearance *AP1* presence was evaluated (32 cycles). Arrows indicate presence and arrowheads absence of *YFP-TCTP1* in roots. Scale bar, 1 cm.

We next assessed mRNA mobility at later developmental stages. Upon prolonged FT induction, when plants entered the reproductive phase and initiated bolting, *YFP-TCTP1* transcript and protein were detected in grafted roots (Figure 2A and 2C). In contrast, plants without FT induction did not show *YFP-TCTP1* transport. To further distinguish developmental phase from plant age, we prolonged the vegetative phase under short-day conditions and delayed flowering. Despite extended growth, *YFP-TCTP1* transcripts were not detected in roots of vegetative plants but were readily detected after transition to flowering under long-day conditions, commencement of floral transition as was confirmed by AP1 expression (Mandel and Yanofsky, 1995, Simon *et al*., 1996) (Figure 2D).

Together, these results demonstrate that restoration of mRNA transport in *dnmt2 nsun2b* mutants depends on the transition to the reproductive phase rather than plant age, indicating that mRNA mobility is controlled by developmental state.

### Vegetative *aly2* and *aly4* Mutants are Deficient in *GRP7* and *TCTP1* Transport

ALYREF proteins are nuclear mRNA export factors that mediate nucleocytoplasmic transport and preferentially bind m^5^C-modified mRNAs in mammalian systems, and presumably also act as m^5^C readers in plants (Zhou *et al*., 2000, Wickramasinghe and Laskey, 2015, Yang *et al*., 2017, Pfaff *et al*., 2018a). An ALY2 ortholog was detected in the *Brassica napus* phloem exudate proteome at very low abundance (4 peptides, 1 unique; see Methods). *A. thaliana* contains four ALY homologs forming two closely related pairs, ALY1/2 and ALY3/4. To test whether ALY proteins contribute to systemic mRNA mobility, we analyzed mRNA transport in *aly2* and *aly4* mutants. We attempted to generate transgenic lines expressing *35S::YFP-TCTP1*, *35S::YFP- HSC70.1* (Figure 1), and as *GRP7* transcript expressed from its own promoter is mobile from shoot to root in juvenile grafts (Figure 3B), we also transferred *GRP7::GRP7-YFP* into the *aly* mutants. Since *35S::YFP-HSC70.1* showed very weak expression in both mutants, it was not further tested. Similarly, as *35S::YFP-TCTP1* showed onset of gene silencing in *aly2* seedlings, these lines could not be used to test *TCTP1* mobility in adult plants. By contrast, *GRP7-YFP* transcripts appeared to be stably produced in both young and mature mutant plants, thus allowing us to use these lines in grafting experiments. *GRP7-YFP* mRNAs produced in shoots were not detected in roots of the grafted plants (Figure 3C). Similarly, *YFP*-*TCTP1* transcripts were detected in *aly4* shoots but not in roots of grafted plants at the vegetative stage (Figure 3C). In line with these observations, the corresponding fusion protein was barely detectable in roots (Figure S2A).

**Figure 3:**
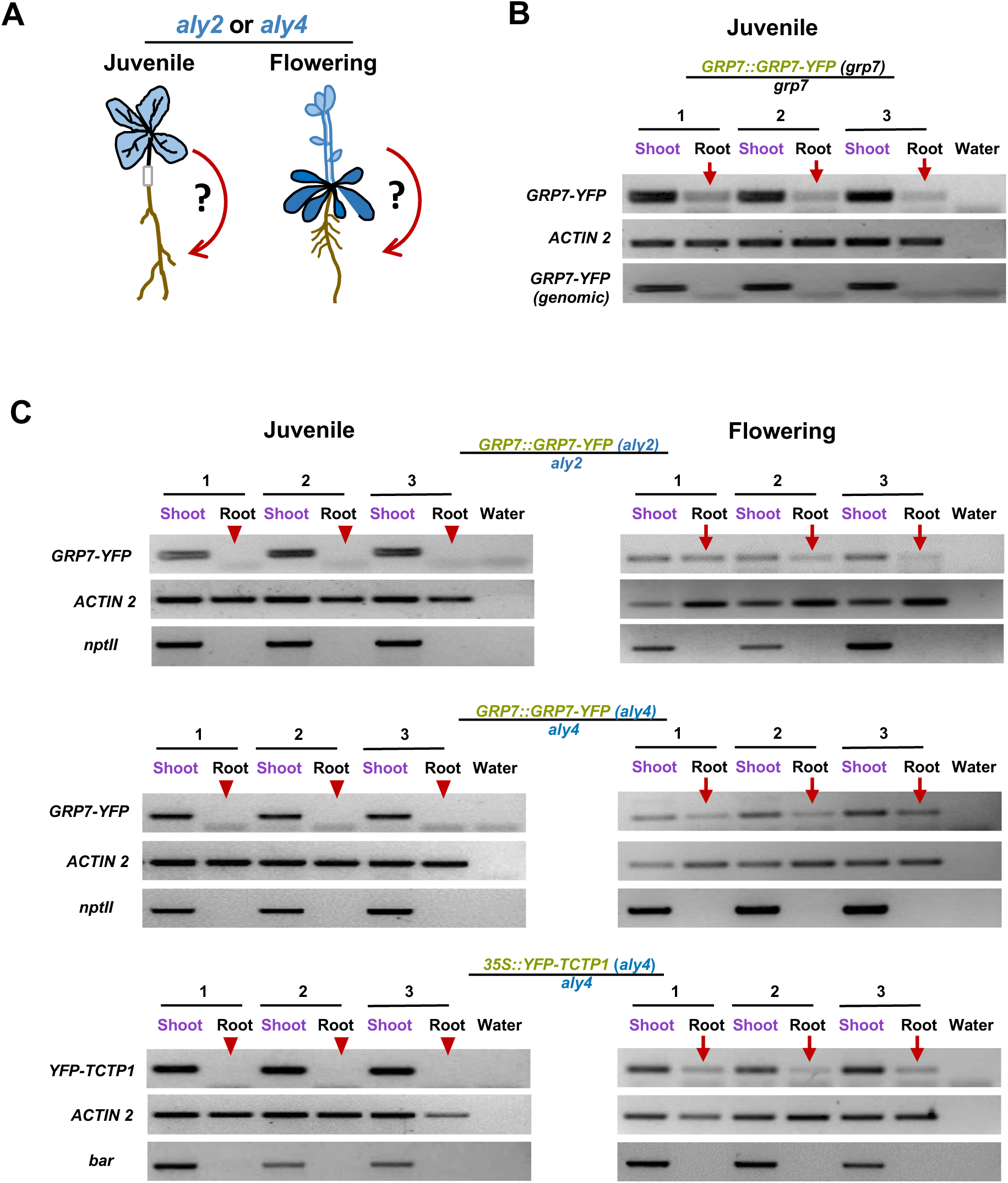
*GRP7* and *TCTP1* mRNA mobility in vegetative and flowering *aly2* and *aly4* mutants. (A) Schematic drawing of transcript mobility analysis performed with hypocotyl-grafted vegetative and flowering *aly2* and *aly4* mutants. (B) RT-PCR assays to detect *GRP7-YFP* mobility in *GRP7-YFP (grp7) / grp7* grafted plants at vegetative stage. (C) RT-PCR assays to detect *GRP7-YFP* and *YFP-TCTP1* mobility in *aly2* and *aly4* mutants at vegetative and flowering stage. Plant tissues from grafted seedlings (n>8) or flowering plants (n=3) were pooled and tested for the presence of the fusion transcripts in grafted tissues. Arrowheads indicate absence, red arrows indicate presence of fusion transcript in roots. For *YFP-TCTP1* and *GRP7-YFP* detection 45 PCR cycles were used. *ACTIN2* primers were used as cDNA quality control (30 cycles). *bar (BastaR)* and *nptII* specific primers were used as a contamination control in the RT-PCR assays (45 cycles). Note that for *GRP7-YFP (grp7)/grp7* samples a *GRP7-YFP* specific genomic PCR control (45 cycles) was performed to confirm that they were not contaminated with transgenic shoot tissue.

We next asked whether the defect in mRNA mobility is developmentally regulated similar to that observed in *dnmt2 nusn2b* mutants. In contrast to vegetative plants, both *GRP7-YFP* and *YFP- TCTP1* transcripts produced in *aly2* or *aly4* shoots were detected in grafted roots at the flowering stage (Figure 3C). Together, these results indicate that ALY2 and ALY4 are involved in systemic mRNA transport during the vegetative growth phase, but are dispensable after the transition to flowering, thus providing further evidence for a developmental regulation of mRNA mobility.

### ALY2 and ALY4 Differentially Bind to m^5^C mRNAs

ALY proteins are mRNA-binding factors involved in nuclear export; however, their associated transcripts *in vivo* remain largely unknown. To address this, we performed RNA immunoprecipitation (RIP) assays followed by sequencing using *ALY2::ALY2-GFP*, *ALY4::ALY4- GFP*, and *35S::3xGFP* control lines (Data S1, Figure S2B, Figure S3). With ALY2, 45 transcripts and with ALY4, 432 transcripts were found significantly enriched (log2FC > 1, P < 0.05) compared to the control (Figure 4A, Data S2). A comparative analysis with published data on Col- 0/Ped-0 graft-mobile transcripts (Thieme *et al*., 2015) revealed a significant enrichment in ALY2 and ALY4 RIP datasets (Figure 4B). For ALY2, 17 out of 45 mRNAs exhibited SNPs between Col-0 and Ped-0, with 10 mRNAs (59%) annotated as mobile. This contrasts with the overall reported transcript mobility rate of 21% detected in Col-0/Ped-0 grafted plants (Thieme *et al*., 2015). Similarly, ALY4 had 196 out of 432 mRNAs with SNPs, of which 75 mRNAs (38%) were reported as Col-0/Ped-0 mobile. Furthermore, a high number of mRNAs (n=18) were common between ALY2- and ALY4-binding transcripts, where 9 had SNPs and of these, 6 mRNAs (66.6%) were reported mobile.

**Figure 4:**
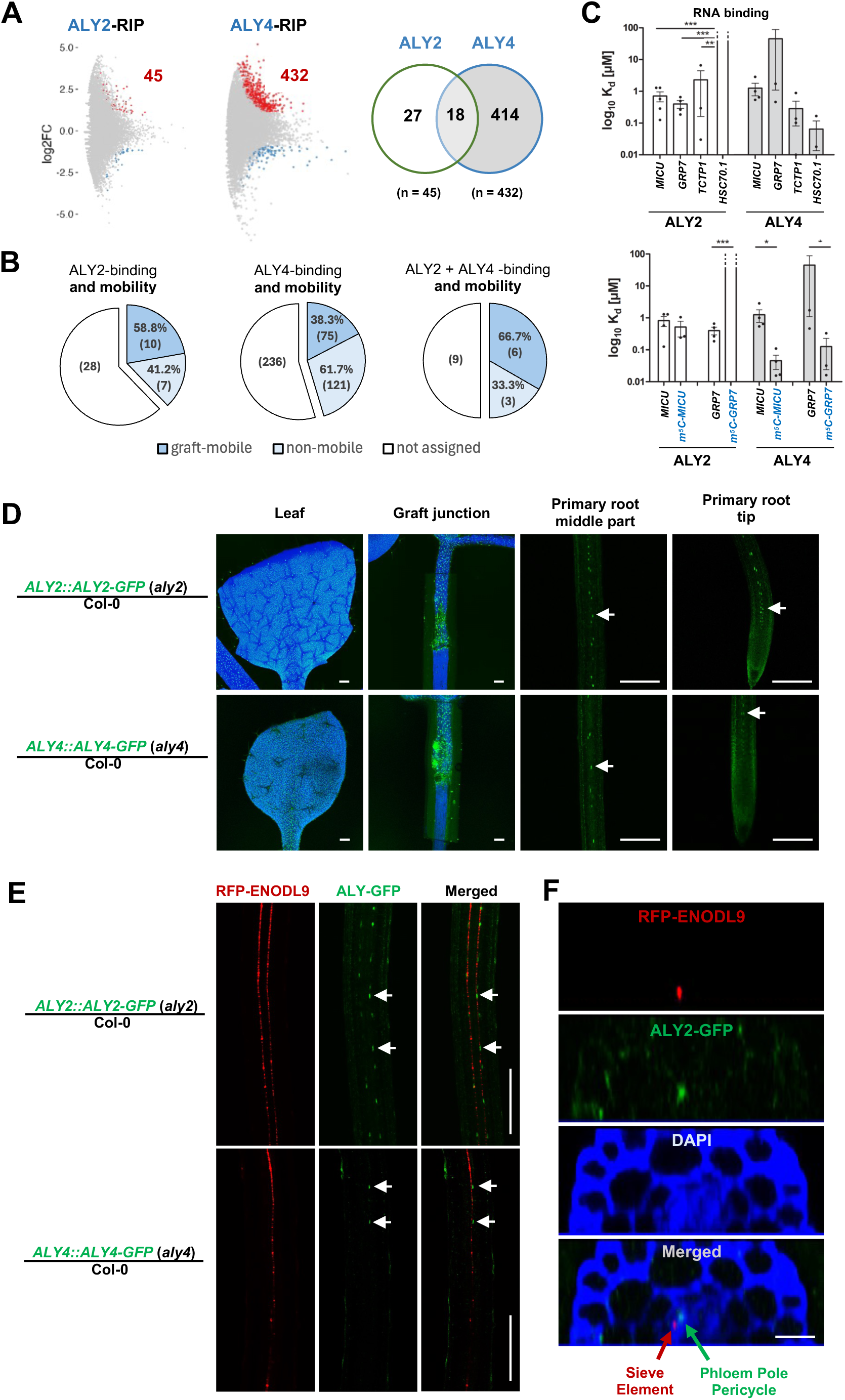
ALY2- and ALY4-binding mRNAs and phloem-mediated shoot-to-root transport. (A) Enriched mRNAs found in ALY2 and ALY4 RNA Immuno-Precipitation (RIP) experiments. (B) Number and fraction of mobile mRNAs found in the ALY2- and ALY4-binding mRNA population. (C) MST measurements on selected mobile mRNAs and *MICU* transcript identified in RIP experiments. *In vitro* synthesized *MICU*, *HSC70.1*, *GRP7* and *TCTP1* RNAs including 5’ and 3’ UTRs were titrated against cell lysates from ALY2-GFP and ALY4-GFP expressing transgenic *Arabidopsis thaliana* plants. Note that ALY4 showed a significantly stronger binding towards *MICU* and *GRP7* m^5^C – methylated mRNAs and ALY2 a significantly weaker affinity to methylated *GRP7*. Statistical significance relative to WT was assessed using one-way ANOVA followed by two-tailed Dunnett’s test with a 95% confidence level. (D) Confocal laser scanning microscope (CLSM) images of GFP fluorescence detected in leaves, graft junctions, and primary roots of grafted vegetative *ALY2::ALY2-GFP /* Col-0 and *ALY4::ALY4-GFP* / Col-0 plants. Blue: auto-fluorescent background of plastids in leaves, arrows indicate presence of fusion proteins in roots. (E) Images of roots of grafted vegetative *ALY2::ALY2-GFP / ENODL9::RFP-ENODL9* and *ALY4::ALY4-GFP* / *ENODL9::RFP-ENODL9* plants. Note that the RFP-ENODL9 (red) sieve element marker is not detected in cells showing ALY2-GFP (green) in nuclei (arrows). Arrows indicate YFP fusion protein presence. Scale bars: 200 µm. (F) Transverse CLSM images of roots from grafted plants. Blue indicates DAPI staining, red arrow indicates RFP presence, green arrow indicates YFP presence. Scale bar: 20 µm.

While this finding is notable, we also acknowledge the possibility that some previously identified candidate mobile mRNAs may have been inaccurately annotated as graft-mobile due to errors in SNP identification and alignment (Liu *et al*., 2019b, Paajanen *et al*., 2025), which could influence the interpretation of the results. Therefore, to be independent of the SNP-based dataset, we compared the two ALY2 and ALY4 RIP datasets with the reported mobile mRNAs moving from Arabidopsis into the phloem-feeding parasitic Cuscuta plant (Thieme *et al*., 2015). Again, the overlap is notably high, with 15 out of 45 for ALY2, and 58 out of 432 for ALY4-binding mRNAs found to be mobile into Cuscuta. Within the set of 18 mRNAs binding to both ALY2 and ALY4, 4 were previously reported to move from Arabidopsis to Cuscuta (Data S2).

To evaluate whether ALY2- and ALY4-binding mRNAs are enriched in m^5^C modifications, we compared the ALY2/4-binding mRNAs with the m^5^C mRNA population of 562 transcripts (Yang *et al*., 2019). Again, a low but significant number of these ALY2-enriched mRNAs (n = 6, p < 1.186e- 04) and ALY4-enriched mRNAs (n = 23, p < 2.317e-06) have been previously annotated as m^5^C-methylated (Yang *et al*., 2019) (Data S2). Supporting the notion that ALY4 preferentially binds to m^5^C-mRNAs, also the proposed m^5^C methylation motif (CC[AUG]CC[AG]) (Yang *et al*., 2019) was found to be significant enriched in the representative gene sequences in ALY4-binding transcripts (158 of 387, 40.8%, p < 2.7e-10), but not in ALY2 targets (14 of 39, 35.9%, p < 0.126). In a more stringent enrichment analysis (log_2_FC > 1, padj < 0.05) of the ALY2 and ALY4 RIP datasets, only one mRNA and 24 mRNAs were found to be significantly enriched in the samples, respectively (Data S2). Here, the transcript encoding the MITOCHONDRIAL CALCIUM UPTAKE (MICU; TAIR# AT4G32060) protein exhibited highly significant enrichment in both RIP samples. MICU regulates mitochondrial calcium uptake (Wang and Teng, 2018, Gutiérrez-Mireles *et al*., 2023) and the transcript was previously assigned as a mobile mRNA candidate in Col-0/Ped-0 grafted plants (Thieme *et al*., 2015). Although *MICU* transcript cannot be found in the stringently analyzed m^5^C mRNA population used in this study (Yang *et al*., 2019), it is present in a less strictly analyzed m^5^C-methylation database (Cui *et al*., 2017).

To further validate the ALY2 and ALY4 interaction with mRNAs and to investigate the potential correlation between ALY2/4-binding mRNAs and m^5^C methylation, we performed MST interaction studies. Here, it should be noted that our attempts to purify ALY2 and ALY4 were hindered by protein aggregation. Furthermore, purified ALY2/4 entities might show different RNA binding characteristics as ALY proteins are part of larger protein complexes (Koroleva *et al*., 2009) and it has been observed that binding characteristics in cell extracts can significantly differ from purified proteins due to cooperative effects of interaction partners present in the cell lysates (Seidel *et al*., 2013, Cao *et al*., 2025, Strauch *et al*., 2025). Thus, we performed Microscale Thermophoresis (MST) RNA–protein interaction assays with cell extracts derived from transgenic *ALY2::ALY2- GFP*, *ALY4::ALY4-GFP*, and *35S::3xGFP* (=control) plants (Ostendorp *et al*., 2022, Fernandes *et al*., 2023, Yang *et al*., 2023b). These extracts were supplemented with *in vitro* produced mRNA and thermophoretic mobility was measured. Both ALY2 and ALY4 bound to three of the tested mRNAs (*MICU*, *GRP7*, *TCTP1*) with reasonable affinities in the nanomolar to micromolar range (Figure 4C). Noteworthy, in contrast to the *MICU* mRNA that was identified in ALY2 and ALY4 RIP assays (see above) and *GRP7* and *TCTP* mRNAs that bind to ALY2 with relative high affinity, the *HSC70.1* mRNA showed poor interaction with ALY2 (estimated kD > 1 mM). ALY4, however, showed strong affinity to *HSC70.1, MICU* and *TCTP1*, and moderate binding to *GRP7* mRNA. ALY2 and ALY4 RNA binding strength was tested with m^5^C-methylated and non-methylated *MICU* and *GRP7* transcripts. m^5^C methylation of *MICU* mRNA did not affect ALY2 RNA binding strength in comparison to non-methylated mRNA, but m^5^C methylation of *GRP7* significantly weakened the interaction with ALY2 compared to non-methylated *GRP7* (p< 0.05; one-way ANOVA). In contrast, ALY4 bound methylated *MICU* and *GRP7* mRNAs with higher affinities than their non-methylated counterparts (Figure 4C). These findings suggest that ALY2 has the capacity to bind to non-methylated and m^5^C methylated mobile transcripts with similar or lower affinity whereas ALY4 binds with higher affinity. However, given that ALY2 and ALY4 were reported to interact with each other in yeast two-hybrid assays (Koroleva *et al*., 2009) both could be necessary *in vivo* to enable delivery of mRNAs from shoot to root despite their different affinities to m^5^C-methylated versus non-methylated mRNAs.

### ALY2 and ALY4 Proteins Move Long-distance from Shoot to Root

ALY proteins have been reported to undergo nuclear-cytoplasmic shuttling in both animals and plants (Zhou *et al*., 2000, Yang *et al*., 2017, Pfaff *et al*., 2018a). However, no evidence besides the low presence of ALY2 in *Brassica napus* phloem exudate harvested from apical stem tissues indicates that the proteins are transported to or via the phloem. To investigate if the two ALY proteins move via the phloem in *A. thaliana*, we grafted wild-type (Col-0) roots with transgenic *ALY2::ALY2-GFP* (*aly2*) or *ALY4::ALY4-GFP* (*aly4*) shoots. Two weeks after grafting both ALY2 and ALY4 proteins fused to GFP were detected at very low levels in nuclei of cells adjoining sieve elements in grafted wild-type roots (Figure 4D to 4F). Notably, both *ALY2-GFP* and *ALY4-GFP* transcripts were not detected in samples of grafted wild-type roots (Figure S4), suggesting that only the fusion proteins and not the transcripts were delivered via the phloem. To identify the root cell type in which ALY fusion proteins appeared, we grafted *ALY2::ALY2-GFP* and *ALY4::ALY4- GFP* shoots with roots harboring the sieve-element specific marker *ENODL9::RFP-ENODL9* (Khan *et al*., 2007). In these roots, GFP-tagged ALY2 appeared in nuclei of phloem pole pericycle (PPP) cells neighboring the red fluorescent RFP-ENODL9-tagged sieve elements (Figure 4E and 4F) indicating that the two ALY proteins exit the sieve elements and accumulate in neighboring PPP cells.

### *aly2/aly4 dnmt2 nsun2b* Triple Mutants Committed to Flowering are Deficient in *GRP7* Transport

Considering the specific lack of *TCTP1* and *HSC70.1* shoot-to-root transport detected in vegetative *dnmt2 nsun2b* (Figure 1) and the lack of *GRP7* mobility in vegetative *aly2* and *aly4* mutants (Figure 3C), we asked whether a more pronounced mRNA transport defect is detected in triple mutants. We generated *aly2 dnmt2 nsun2b* and *aly4 dnmt2 nsun2b* triple mutants by crossing, and grafted these with *grp7* mutant roots to assay endogenous *GRP7* transcript mobility in flowering plants. In contrast to grafted *dnmt2 nsun2b* double mutants (Figure S5) and *aly2* and *aly4* single mutants (Figure 3C), the *GRP7* transcript did not move from shoot to root in the triple mutants committed to flowering (Figure 5A). These findings indicate a genetic interaction between the m^5^C methyltransferase and the two ALYs and supports the notion that ALY2 and ALY4 act downstream of or in concert with the methyltransferases to facilitate transcript transport in flowering plants.

**Figure 5:**
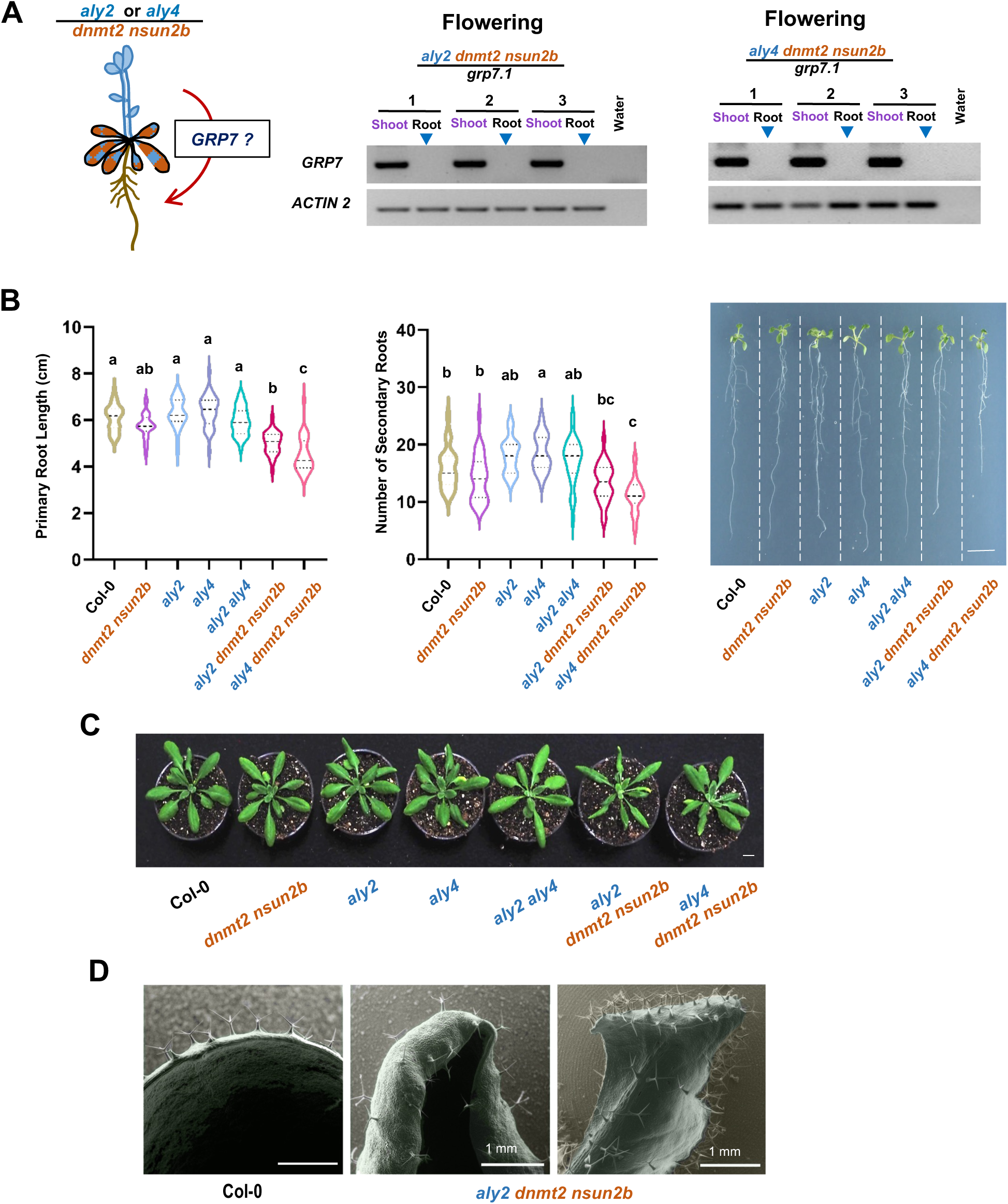
Plant growth and mRNA mobility in *aly dnmt2 nsun2b* triple mutants. (A) Schematic drawing and PCR assays used to detect endogenous produced *GRP7* transcript transport to *grp7* roots from grafted scions lacking methyltransferase activity and ALY2 or ALY4 expression. Note that after 45 PCR cycles no *GRP7* transcript was detected in root RNA samples (arrowheads) from grafted flowering *aly2 dnmt2 nsun2b / grp7* or *aly4 dnmt2 nsun2b / grp7* plants. (B) Primary root length and lateral root number of 10-day-old wild-type (Col-0), *dnmt2 nsun2b, aly2, aly4, aly2 dnmt2 nsun2b* and *aly4 dnmt2 nsun2b* mutant seedlings. Scale bar: 1 cm, n = 30; significance was evaluated using one-way ANOVA (α = 0.05) followed by multiple comparisons of means using Tukey’s HSD test at the 0.05 significance level. (C) Representative images of 26-day-old wild-type (Col-0) and mutant plants grown on soil. Note the appearance of aberrant shaped rosette leaves on *aly2 dnmt2 nsun2b* mutants. Scale bar: 1 cm, n = 21. (D) Representative Scanning electron micrographs (SEM) of rosette leaves of Col-0 and *aly2 dnmt2 nsun2b*.

Next, we asked whether root growth or flowering time is changed in *aly2 dnmt2 nsun2b* and *aly4 dnmt2 nsun2b* triple mutants compared to *aly2 aly4* and *dnmt2 nsun2b* double mutants. Primary root length and the number of lateral roots were significantly reduced in *aly2 dnmt2 nsun2b* and *aly4 dnmt2 nsun2b* triple mutants compared with the corresponding single or double mutants (Figure 5B). Similar to *dnmt2 nsun2b*, the *aly2 aly4* double mutants showed an early flowering phenotype which was not significantly different from that of the triple mutants (Figure S6). Notably, *aly2 dnmt2 nsun2b* triple mutants developed downward-curled rosette leaves (Figure 5C and 5D) resembling the phenotype reported for the ALY2-interacting HAPLESS 1 / MAGO protein upon knockdown (Park *et al*., 2009). This leaf phenotype was not observed in any of the corresponding single or double mutants suggesting a functional linkage between m^5^C RNA methylation and ALY function.

To find further support for a potential link between mRNA m^5^C methylation and nuclear export factors we asked whether ALY2/4-related ALY1 (AT5G55950) / ALY3 (AT1G66260) and NSUN2B / DNMT2-related methyltransferases were induced in the *aly2, aly4* or *nsun2b dnmt2* mutants potentially facilitating mRNA mobility during flowering. The closest paralog of *NSUN2B* (also named *TRM4B)* is *TRM4A* (AT4G40000), whereas *DNMT2* does not appear to have a predicted paralog in the genome that could compensate for its loss. None of methyltransferase-related genes nor *ALY1* and *ALY3* were significantly differentially expressed in *aly2, aly4* or *dnmt2 nsun2b* vegetative plants, or *dnmt2 nsun2b* mutants committed to flowering (Table S1, Data S3). Consistently, the expression of mobile transcripts, including *GRP7*, *HSC70.1*, and *TCTP1*, were not significantly changed across genotypes and developmental stages, except for the reduced *TCTP1* and *HSC70.1* transcript levels detected in the *aly2* mutant. Thus, the observed change of mobility in vegetative vs flowering committed mutants seems not to be a consequence of induced gene expression of the methyltransferases and/or ALY-related export factors potentially compensating for the loss of the respective family member.

## DISCUSSION

Our data demonstrate that in vegetative *dnmt2 nsun2b* mutants, the transport of *TCTP1* and *HSC70.1* mRNAs from shoot to root is severely impaired, consistent with prior reports (Yang *et al*., 2019, Yang *et al*., 2023b). However, in plants committed to flowering, this transport is fully restored, even in the absence of functional m^5^C methyltransferases. This suggests that the m^5^C modification, while essential in juveniles, becomes dispensable in adults. The restoration of mobility is not due to compensatory upregulation of *DNMT2* or *NSUN2B*, nor is it driven by increased expression of the mobile transcripts themselves. Instead, it reflects a developmental reprogramming of the mRNA transport machinery. This phenomenon is not simply a consequence of aging, as prolonged vegetative growth under short-day conditions fails to restore transport, whereas floral transition under long-day conditions does. Furthermore, induction of *FT* expression alone - sufficient to initiate flowering - does not restore mobility, indicating that the transition to the reproductive phase, rather than florigen signaling per se, is the key trigger. This implies that *TCTP1* and *HSC70.1* can use alternative mechanisms - possibly involving different RNA-binding proteins or structural motifs - to bypass the need for m^5^C methylation in the reproductive phase.

### ALY2 and ALY4 as Developmentally Regulated Facilitators of mRNA Export

In search of factors interacting with m^5^C modified mobile mRNAs we identified ALY2 and ALY4 – members of the ALYREF protein family shown to bind preferentially to m^5^C-methylated mRNAs (Yang *et al*., 2017, Chen *et al*., 2019, Yang *et al*., 2019) - that have long been implicated in nuclear mRNA export via the TREX complex, linking transcription, splicing, and export. Our results now extend this role to systemic mRNA transport, revealing that ALY2 and ALY4 are critical for the delivery of mobile mRNAs such as *TCTP1* and *GRP7* in vegetative plants. In *aly2* and *aly4* single mutants, shoot-to-root transport of these transcripts is abolished during vegetative growth, but is restored in flowering plants - mirroring the behavior of *dnmt2 nsun2b* mutants. This parallel developmental regulation strongly suggests that ALY2 and ALY4 function in a shared pathway with m^5^C methyltransferases during juvenile development.

These findings imply - supported by the observed lack of induced expression in the methylation mutant (Table S1, Data S3) - that alternative factors besides methylation or the ALYs are sufficient to facilitate mRNA transport at later developmental stages. Adding to the complexity of the mRNA transport system, *GRP7* transcript transport appeared to be affected by lack of either ALY2 or ALY4 in vegetative but not in flowering plants. Notably, *GRP7* mobility was completely abolished only in *dnmt2 nsun2b aly2* and *dnmt2 nsun2b aly4* triple mutants. Given that *GRP7* expression was not changed in the *dnmt2 nsun2b* and the *aly2* and *aly4* mutants (Table S1, Data S3), this suggests that the lack of transport of the endogenous produced *GRP7* transcript is not caused by decreased abundance of its transcript. The combined activities of the m^5^C writers and the ALY nuclear export factors are necessary to facilitate *GRP7* transport in flowering plants. This notion is supported by the curly leaf phenotype detected exclusively in *aly2 dnmt2 nsun2b* triple mutants suggesting genetic interaction (Figure 5).

Based on RIP assays, both nuclear RNA export factors seemed to bind preferentially, but not exclusively, to mobile mRNAs with an m^5^C methylation motif (Figure 4A and B). However, MST measurements indicated that ALY4 binds preferentially to methylated and ALY2 to non-methylated mRNAs displaying different selectivity. In line with a potential m^5^C reader function, only ALY4 showed significantly higher affinity to m^5^C-methylated *MICU* and *GRP7* mobile mRNAs, with *MICU* mRNA enriched in both ALY2 and ALY4 RIP assays. ALY2 showed no difference for *MICU* RNA, methylated or not, but the affinity to *in vitro* methylated *GRP7* RNA appeared strongly reduced (Figure 4A and C). Seemingly, these findings contradict the observation that *GRP7* transport is abolished in flowering *aly2/4 dnmt2 nsun2b* triple, but not in vegetative and *dnmt2 nsun2b* double or in *aly2* and *aly4* single mutants committed to flowering. The surprising finding that *GRP7* mobility can be independent of m^5^C activity in juvenile plants (Figure 6A) might be explained by *GRP7* harboring a mRNA transport motif that is formed without assistance of m^5^C modifications. That *GRP7* transport is not affected in *aly2* and *aly4* single mutants but in *dnmt2 nsun2b aly2/4* triple mutants committed to flowering could be explained by *GRP7* m^5^C methylation and dimer formation of ALY2 and ALY4 (Koroleva *et al*., 2009, Arabidopsis Interactome Mapping, 2011) and by their presence in the same nuclear export complex (Pfaff *et al*., 2018a). The lack of ALY2 may affect the preferential binding of ALY4 to m^5^C-methylated transcripts and *vice versa*, and, by this means impair *GRP7* mRNA transport in vegetative plants. However, in plants committed to flowering both the nuclear export factors as well as m^5^C modifications appear to be essential for efficient transcript delivery. In other words, it seems that, depending on their different binding affinity towards m^5^C-modified or non-modified mobile mRNAs, such as *TCTP1* or *GRP*7, respectively, they act together deciding which mRNAs can be exported and enter the phloem long-distance transport pathway towards roots.

**Figure 6:**
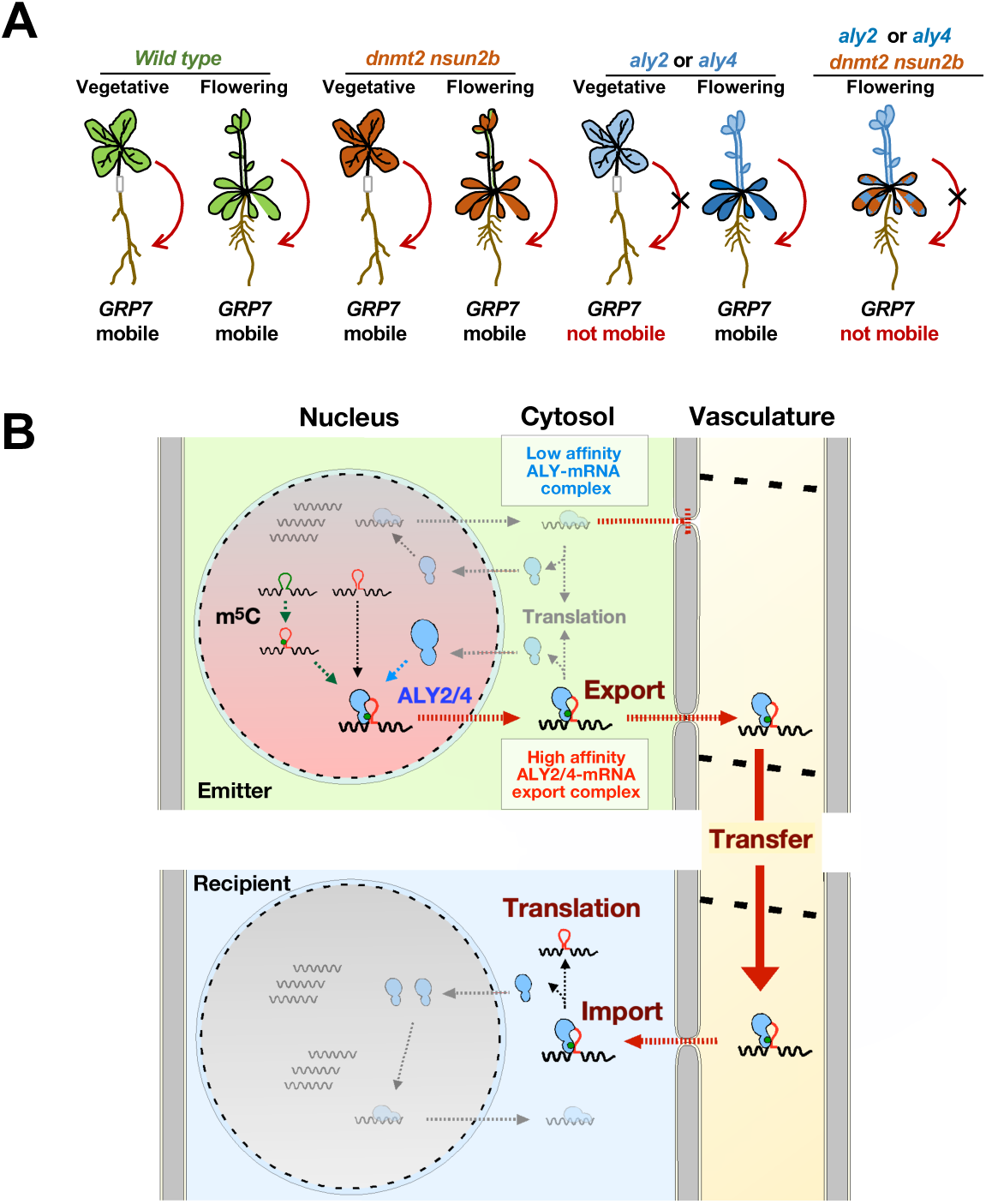
Schematic models illustrating ALY2/4 function in the regulation of mRNA transport. (A) Schematic drawing showing *GRP7* transcript transport from scion to rootstock in grafted wild-type plants, and *aly2/4* and *dnmt2 nsun2b* mutant plants. (B) Schematic drawing of ALYREF function in mediating mRNA transport. ALY2 and ALY4 proteins (blue) move as large RNP complexes out of the nucleus into the cytosol. Here, mRNAs bound to ALY with weak affinity (e.g., mRNAs lacking a mobility motif and/or the m^5^C modification) are released for translation. Transcripts with m^5^C modifications and/or a mobility motif (red) are bound to ALYs with high affinity and may move as RNA-protein complexes via plasmodesmata from companion cells to sieve elements (vasculature) and, consequently, to roots. Here, ALY2/4 RNPs are unloaded into recipient cells, such as the phloem pole pericycle cells. Prior to their nuclear import, the delivered mRNA cargoes are either released from the ALY-RNA complexes for translation in recipient cells or are further transported to more distant cells.

### ALY Proteins Are Themselves Phloem-Mobile: A Novel Mechanism for Systemic mRNA Delivery

To identify factors that discriminate between m^5^C methylated and non-methylated mRNAs and may mediate shoot to root transcript delivery, we identified the RNA binding proteins ALY2 and ALY4 as phloem mobile proteins (Figure 4C to E). The phloem pole pericycle (PPP) is known to serve as the repository for phloem-unloaded macromolecules (Ross-Elliott *et al*., 2017), including *TCTP1* (Yang *et al*., 2019). One of the most striking findings of this study is the detection of ALY2-GFP and ALY4-GFP proteins in the nuclei of PPP cells in grafted wild-type roots. This demonstrates that ALY proteins are not only involved in nuclear export but are themselves transported long-distance via the phloem. The absence of *ALY2* and *ALY4* transcripts in grafted roots confirms that ALY protein presence in the roots is not due to local transcription of delivered mRNAs but rather to transport of the protein itself. This is a crucial observation: ALY proteins, traditionally viewed as nuclear factors, are delivered to recipient tissues where they may facilitate the import and/or processing of mobile mRNAs.

Consistent with function in the cytosol and mRNA long-distance transport, ALYs also seem to play a role in viral RNA transfer and infection spread. Polerovirus Turnip Yellow Virus (TuYV) coat protein interacts with the four ALY proteins leading to a change in their intranuclear distribution and altered RNA export (Kiervel *et al*., 2024). Export of Cauliflower Mosaic Virus (CaMV) polycistronic viral RNA involves numerous protein partners, among which ALY proteins play a prominent role (Ehrnsberger *et al*., 2019). Deficiency in ALY factors resulted in plants that were partially resistant against CaMV infection due to impaired viral RNA export from the nucleus to the cytoplasm (Kubina *et al*., 2021). ALY proteins can also be retained in the cytosol by interacting viral factors as shown with Tomato bushy Stunt Virus (TbSV). All four ALYs interact with the TbSV P19 factor to interfere with its ability to suppress RNA silencing of viral RNA (Uhrig *et al*., 2004, Canto *et al*., 2006).

The combined findings support a model (Figure 6B) in which ALY proteins act as “molecular shuttles” that carry mobile mRNAs from the shoot to the root. In this model, ALY2 and ALY4 bind to mobile mRNAs in the shoot, facilitate their export from the nucleus, and then accompany them through the phloem. Upon reaching the root, they may assist in the release of the mRNA from the phloem into recipient cells - possibly via plasmodesmata that apparently contain nuclear pore complex (NPC) components (Schladt *et al*., 2025). This would explain how mRNAs can traverse the phloem and enter recipient cells without being degraded or trapped in the vascular system. After phloem unloading of the mobile complex to the recipient phloem pole pericycle cells (Figure 4D to F) the delivered mRNA cargo could be released for translation, or may be further transported to more distant cells.

The mobility of ALY proteins also provides a potential explanation for the developmental switch in mRNA transport. In vegetative plants, ALY2 and ALY4 may be required to bind and escort mRNAs through the phloem, possibly in conjunction with m^5^C modifications that enhance their stability or recognition. In flowering plants, alternative export factors or RNA-binding proteins may take over this role, allowing transport to proceed independently of m^5^C methylation and ALY2/4 function - except when both are simultaneously compromised. These findings redefine our understanding of plant systemic signaling by adding another layer of complexity. Although our data show that ALY2/4 and m^5^C methylation are both involved in mRNA delivery, we still have to identify the molecular factors determine the developmental dependency of mRNA transport.

## MATERIALS AND METHODS

### Plant material and expression constructs

The Col-0 mutants *dnmt2* (SALK_136635) *nsun2b* (SAIL_667_D03) were provided by Dr. lain Searle laboratory, Adelaide) (Burgess *et al*., 2015), and *grp7.1* (SALK_039556.21.25.x) were provided by Prof. Dorothee Staiger (University of Bielefeld, Germany). *Arabidopsis thaliana aly2* (wiscDsLox461-464N10), *aly4* (GK-497B06), *ALY2::ALY2-GFP* (*aly2*) and *ALY4::ALY4-GFP* (*aly4*) were provided by Prof. Klaus Grasser (University of Regensburg, Germany) (Pfaff *et al*., 2018b), *pER22-SUC2:FT-GFP ft-10* was provided by Prof. Hao Yu (National University of Singapore, Singapore) (Liu *et al*., 2019a). *Agrobacterium tumefaciens* AGL1 strain containing *35S::YFP-TCTP1* expressing binary constructs and *Arabidopsis thaliana 35S::YFP-TCTP1* (Col- 0), *35S::YFP-TCTP1* (*dnmt2 nsun2b*), *35S::HSC70.1-GFP* (Col-0) and *35S::HSC70.1-GFP* (*dnmt2 nsun2b,* Col-0) lines were previously described (Yang *et al*., 2019). To create the *GRP7::GRP7-YFP* expression vector, the 2233 bp *GRP7* promoter (upstream of start codon), the *GRP7* CDS lacking the stop codon, and *YFP* with the *GRP7* 3’UTR (476 bp downstream of stop codon) were PCR amplified (for PCR primers see Data S5) to introduce restriction sites for ligation into SalI / XhoI digested pENTR4 gateway vector. Then, the construct was transferred via gateway reaction into the pGWB401 binary vector. The *GRP7::GRP7-YFP* and *35S::YFP-TCTP1* binary vectors were used to produce the transgenic *Arabidopsis thaliana* (Col-0) *GRP7::GRP7- YFP* (*grp7*), *35S::YFP-TCTP1* (*aly2*), *35S::YFP-TCTP1* (*aly4*), *GRP7::GRP7-YFP* (*aly2*) and *GRP7::GRP7-YFP* (*aly4*) lines by floral dipping (Davis *et al*., 2009).

### Arabidopsis hypocotyl grafting

Sterilized *Arabidopsis thaliana* Col-0 seeds were stored at 4 °C for two days, then germinated on plates (1/2 MS, 1% sucrose, 1% agar) placed vertically under short day (SD) conditions (9 h light / 16 h dark, 21 °C, light intensity: 170 μE·m^−2^·s^−1^) in a Percival growth chamber. 6-7 days old seedlings (after germination) were cut in the upper half of the hypocotyl with a vertical stroke using a surgical scalpel, and silicon micro tubes (VWR) with 0.3 mm diameter were used to support graft junctions. Grafted plants were grown vertically on new plants as above. Adventitious roots from the scions started to be removed one week after grafting.

### Plant growth conditions and harvesting

If not otherwise mentioned, sterilized seeds and grafted plants were grown on half-strength (1/2) Murashige and Skoog (MS) medium supplemented with 1% (w/v) sucrose and solidified with 1% (w/v) microagar (Duchefa Biochemie). Plates with grafted seedlings were placed vertically in Percival growth chambers set to short day (SD) conditions (9 h light / 16 h dark, 21 °C, light intensity: 170 μE·m^−2^·s^−1^). Vegetative plants were harvested 12-14 days after grafting. For producing flowering (flowering) plants, grafted plants (12-14 days after grafting) were transferred to full nutrient liquid cultures with no sucrose (1 mM KNO_3_, 0.5 mM NH_4_NO_3_, 1.5 mM KH_2_PO4/K_2_HPO_4_ pH 5.7–5.8, 2 mM CaCl_2_, 0.5 mM MgSO_4_, 1 mM K_2_SO_4_, 1.5 mM MES, 2 μM Na_2_FeEDTA, 30 μM H_3_BO_3_, 7 μM MnSO_4_, 0.5 μM ZnSO_4_, 0.3 μM CuSO_4_, 0.2 μM, NiCl_2_, 0.15 μM HMoO_4_, 10 nM CoCl_2_), and grown in growth chambers set to neutral day conditions (12 h light / 12 h dark, 22 °C, light intensity: 170 μE·m^−2^·s^−1^), shaking at 70 rcf. The plants were transferred to fresh liquid medium every 2-3 days until harvest.

For flowering and mobility experiments shown in Figure 2 grafted vegetative plants were first grown on ½ MS medium supplemented with 1% sucrose and solidified with 1% (w/v) microagar (Duchefa Biochemie) (shown in Figure 2b). Then, grafted plants were transferred to soil around 10 days after grafting. After transferring to soil, grafted plants were grown in Percival growth chambers under long day (LD) conditions (16 h light / 8 h dark; day 22 °C / night 19 °C; light intensity: 170 μE·m^−2^·s^−1^; relative humidity: 60%) (shown in Figure 2c); or were grown in Percival growth chambers under SD conditions (8 h light / 16 h dark; 16 °C; light intensity: 100 μE·m^−2^·s^−1^; relative humidity: 60%), then harvested after 40 days, or transferred to LD conditions (16 h light / 8 h dark; day 22 °C / night 19 °C; light intensity: 170 μE·m^−2^·s^−1^; relative humidity: 60%)(Figure 2d).

To produce seedlings for RIP experiments (see methods below), sterilized seeds were grown on ½ MS medium supplemented without sucrose and solidified with 1% (w/v) agar. Plates were placed vertically in Percival growth chambers set to LD conditions (16 h light / 8 h dark, 22 °C, light intensity: 170 μE·m^−2^·s^−1^). The seedlings were harvested for RIP experiment 10 days after germination.

### β-Estradiol induction of *pER22-SUC2*::*FT-GFP*

β-Estradiol (10 µM) was applied on rosette leaves, in young grafts grown on plates, two days after treatment, shoots and roots were harvested and analyzed. For grafted plants grown on soil, rosette leaves were treated every day starting from 7 days after transfer to soil. Shoots and roots were harvested for analysis around 14 days after β-Estradiol treatment when the plants bolted. The induced leaves were dissected and examined by Leica SP8 confocal laser scanning microscope (CLSM) for the presence of GFP-FT expression in the vasculature.

### ALY protein identification in *Brassica napus* phloem exudate and leaves by LC-MS/MS

For the identification of *B. napus* phloem ALY-related proteins, phloem sap was collected from flowering plants and precipitated to enrich proteins as described (Ostendorp *et al*., 2017). In short, the protein pellet was resuspended in the same volume of extraction buffer (6□M urea, 2□M thiourea, 15□mM DTT, 2% CHAPS) as the original volume of collected phloem exudate. The protein solution was sonicated for 10□minutes in a sonication bath, followed by 30□minutes of incubation on an orbital shaker (100□rpm) at room temperature. Solubilized proteins were centrifuged at 10,000□×□g for 5□min to remove non-solubilized aggregates, and the protein concentration was determined in the collected supernatant.

For protein analysis of leaf tissue from flowering plants, the leaves were harvested, grinded in liquid N_2_ and the powder was incubated in extraction buffer (6□M urea, 2□M thiourea, 15□mM DTT, 2% CHAPS) added to a final protein concentration of 2□µg/µl. 25□µg of phloem sap and 50□µg of leaf tissue protein extract were then digested in solution using a Trypsin/Lys-C mixture (Mass Spec Grade, Promega, Madison, WI, USA) according to the instruction manual. After the digestion, the samples were desalted using C18-stage tips as described (Rappsilber *et al*., 2007). After the elution of the digested and desalted peptides from C18-stage tips, the samples were concentrated to near dryness in a SpeedVac and the peptide mixtures were reconstituted in 30 µl resuspension buffer (5% acetonitrile, 0.1% formic acid in water). 5□µl of this peptide mix was injected onto an Acclaim PepMap RSLC HPLC column (75□µm□×□15□cm, Thermo Scientific) connected to the EASY-nLC 1000 system (Thermo Scientific). The eluting peptides were analyzed on a Q Exactive Plus (Thermo Scientific, Bremen, Germany) high-resolution mass spectrometer. Peptides were separated using a binary buffer system of 0.1% formic acid in water (Buffer A) and 60% acetonitrile containing 0.1% formic (Buffer B). The flow rate was adjusted to 300□nl/minute. Peptides were eluted with a linear gradient from 0–40% buffer B for 50□min followed by a linear gradient between 40–80% buffer B for additional□30 min. Peptides were analyzed in the mass spectrometer using one full scan (300–1600 m/z, R□=□70,000 at 200□m/z), followed by up to fifteen data-dependent MS/MS scans (Top 15 approach) with higher-energy collisional dissociation (HCD) at a resolution of 17,500 at 200□m/z. Dynamic exclusion was set to 15□s. Raw data were processed using the Progenesis QI for proteomics (Progenesis QI for Proteomics Version 3.0, Nonlinear Dynamics, Newcastle, UK) software and the protein sequences of all identified ALY-related proteins from *B. napus* as described (Ostendorp *et al*., 2017). Protein identifications were filtered with a false discovery rate better than 1%, at least two peptides, one unique peptide and a score of 50. For the ALY orthologous *B. napus* protein (BnaA03g54780D), 4 peptides and 1 unique peptide were found to be present in the phloem exudate sample.

### RNA isolation

Grafted shoots and roots were separately harvested in 1.5 ml tubes containing metal beads and frozen in liquid nitrogen. After grinding, the broken tissue was supplemented with 0.75 ml Trizol Reagent (Invitrogen), homogenised (20s vortex), and incubated for 5 minutes at room temperature. After chloroform extraction (0.2 mL per 0.75 mL of TRIzol Reagent), samples were vortexed and incubated for 5 minutes at RT and centrifuged for 15 minutes (12□000 g at 4°C). To digest DNA, 1 μl DNaseI (Thermo Scientific) and 0.5 μl RNasin (Promega) were added to the upper aqueous phase, and incubated for 15 minutes at RT. After 0.1 ml chloroform extraction, upper RNA phase was transferred to 1.5 mL RNase-free tubes and 1 volume isopropanol and 1/10 volume 3 M NaOAc (pH 5.2) were added to precipitate total RNA, invert several times, incubated overnight at −20°C. After centrifugation for 30□minutes at 4°C and the RNA pellets were washed two times with 1□ml ice-cold 80% ethanol, one time with 99% ethanol, dissolved in 10–25□μl DEPC H_2_O, and stored at −80°C.

### Differential expression analysis of vegetative seedings

Total RNA samples (n=3) from pools (>5 plates with > 50 seedlings for each pool) of 10-day-old Col-0 and *dnmt2 nsun2b* seedlings grown on ½ MS plates supplemented with 1% sucrose were harvested end of day and submitted to Illumina cDNA library production and sequenced (±50 million reads paired-end RNAseq; BGI, Shenzhen, China).

The readouts were mapped to the Arabidopsis genome TAIR10 version GCF_000001735.4 (https://www.ncbi.nlm.nih.gov/datasets/genome/GCF_000001735.3/) with STAR (v2.7.9a)(Dobin *et al*., 2013). The counts of transcripts are then quantified by featureCounts (v2.0.6)(Liao *et al*., 2014). Differentially expressed genes were identified using DESeq2 (Love *et al*., 2014) by comparing 10-day-old *dnmt2 nsun2b* samples against control samples (Col-0). Genes exhibiting a log2-fold change greater than 1 were classified as up-regulated, while those with a change less than -1 were regarded as down-regulated. Both groups of genes are considered significantly dysregulated with p-value blow 0.05. Enrichment analysis for Gene Ontology (GO) and KEGG pathways of DEGs was performed using the R package clusterProfiler (Wu *et al*., 2021). The reference genes for this analysis were those with a TPM greater than 1 in either control or *dnmt2 nsun2b* samples. The most significant 10 pathways with smallest adjusted p-values are presented in Figure S6b.

### Microscopy

To detect the presence of RFP-ENODL9, YFP-TCTP1, FT-GFP, ALY2-GFP and ALY4-GFP fusion proteins in grafted Arabidopsis plants, either a Leica SP5 or SP8 confocal laser scanning microscope (CLSM) was used, XYZ and YXZ stack images were assembled and processed using the Fiji software package as described (Yang *et al*., 2019).

### RNA immunoprecipitation (RIP)

Transgenic seeds were vertically grown on plates under SD conditions for 10 days (see methods above on growth conditions), then subjected to irradiation with 254 nm UV light at a dose of 500 mJ / cm2 (UVP CL-1000 UV crosslinker) on ice and frozen in liquid nitrogen immediately after crosslinking. For each replicate, 3 g sample was ground in liquid nitrogen and protein was extracted with 4.5 mL lysis buffer (150 mM NaCl, 50 mM Tris-HCL pH7.5, 4 mM MgCl_2_, 1% SDS, 0.25% Deoxycholic acid sodium salt, 0.25% Igepal, cOmplete™ Protease Inhibitor Cocktail (Roche, 1 tablet per 50 mL), 0.5 mM DTT, 1 mM PMSF, 100 U/mL RNasin® Ribonuclease Inhibitor (Promega)), incubated at 4°C and 1400 rpm for a few seconds, until solution becomes homogenous. After centrifugation (10 minutes, 16000 g, 4°C), supernatant was filtered by 0.45 µm filter and transferred to a new tube supplemented with 25 μL equilibrated Binding Control Magnetic beads (Chromotek) lacking GFP-Trap® antibody for preincubation, mixed by gentle end-over-end mixing for 30 minutes at 4°C. The magnetic beads were magnetically separated and discarded and the cleaned cell lysates were transferred to new tubes and incubated with 25 µL equilibrated of GFP-Trap® M beads (Chromotek) and rotated (10 rpm) for 4 h at 4°C. Supernatant was removed from magnetically separated beads, and the beads were washed one time with 8 mL ice-cold wash buffer (500 mM NaCl, 50 mM Tris-HCL pH7.5, 4 mM MgCl_2_, 1% SDS, 0.5% Deoxycholic acid sodium salt, 2M Urea), four times with 4 mL ice-cold wash buffer, and three times with ice-cold PNK buffer (20 mM Tris-HCl, pH 7.4, 10 mM MgCl_2_, 0.2% Tween 20). Then RNA extraction was performed using the Trizol protocol (see above RNA extraction) and submitted to RNA sequencing and/or RT-PCR.

### RNA sequencing and analysis of RIP

Total RNA from three replicated RNA immunoprecipitation (RIP) samples from *35S::3xGFP*, *ALY2-GFP and ALY4-GFP* transgenics were submitted to Illumina cDNA library production and sequencing (±20 million reads paired-end RNAseq; BGI, Shenzhen, China). The readouts were mapped to the Arabidopsis genome TAIR10 version GCF_000001735.4 (https://www.ncbi.nlm.nih.gov/datasets/genome/GCF_000001735.3/) with STAR (v2.7.9a) (Dobin *et al*., 2013). The counts of transcripts are then quantified by featureCounts (v2.0.6) (Liao *et al*., 2014). The differentially binding transcripts were detected by DESeq2(Love *et al*., 2014) by comparing the ALY2-GFP or ALY4-GFP sample versus *35S::3xGFP* control samples. The transcripts with log2-fold changes (p < 0.05, or padj < 0.05 in a stringent analysis) are selected as enriched transcripts. For motif enrichment analysis, we utilized ‘SEA’ (Bailey and Grant, 2021). Representative gene sequences were analyzed against shuffled sequences preserving 3-mer frequencies as controls, to validate the enrichment and significance of the m^5^C methylation motif.

### *In vitro* RNA transcription

RNAs including 5’ and 3’ UTRs were synthesized *in vitro* using T7 RNA polymerase kit (Promega). cDNA from A*. thaliana* Col-0 were used to amplify single transcripts for *HSC70.1, MICU*, *GRP7* and *TCTP1*. 5 to 10 pmol of purified PCR fragments were used per 100 µl transcription reaction. Reactions were incubated at 37°C for 3 h followed by a DNA digestion for 30 min. Subsequently, all RNAs were purified using the RNA clean & concentrator-25 kit (ZymoResearch). RNA purity and integrity were checked using Nanodrop One and BioAnalyzer PicoChip (Agilent).

### Microscale thermophoresis on ALY2 and ALY4 cell lysates

Lysates were prepared according to (Yang *et al*., 2023b). In brief, a single 3 -weeks-old plant grown on ½ MS agar plate was frozen in liquid nitrogen and subsequently ground to a fine powder. 200 µl of 2x MST buffer (100 mM HEPES-NaOH pH 7.5, 300 mM NaCl, 20 mM MgCl_2_, 0.2 % NP-40, 2 mM AEBSF, 2 mM PMSF, 0.5 U/µl RiboLock RNase inhibitor) was added and incubated for 15 min on ice. Lysates were centrifuged at 21000 x g for 10 min. Subsequently, the cell extract was further diluted with 2x MST buffer until fluorescence reads between 1600 and 400 counts were achieved. RNAs were set to an initial concentration of max 2 µM and 1:1 dilution series were made with water. Finally, 5 µl of cell extract was added to 5 µl of each RNA dilution and incubated for 5 min at room temperature. Samples were measured in standard capillaries on a Monolith NT.115 (Nanotemper) and analysed using MO:Analysis software and GraphPad Prism 5. Binding was regarded as present as soon as a signal-to-noise ratio and response amplitude larger than 5 was achieved.

## DATA AVAILABILITY STATEMENT

The sequencing data are available at the NCBI Sequencing Read Archive (SRA), BioProject ID PRJNA1082496.

## ACKNOWLEDGEMENTS

We thank Melissa Tomkins and Richard Morris (JIC, UK), Julia Kehr (University of Hamburg, Germany) and Ralph Bock for helpful comments on the manuscript text and data analysis. Dana Schindelasch and Cindy Hauptvogel (MPI-MPP) for excellent technical support. We thank Dorothee Staiger (University of. Bielefeld, Germany) for providing *grp7.1* mutant seeds and discussions on the RIP experiment, Klaus Grasser (University of Regensburg, Germany) for generously providing *aly2*, *aly4*, *ALY2::ALY2-GFP* (*aly2*) and *ALY4::ALY4-GFP* (*aly4*) lines and helpful comments on the manuscript, and Hao Yu (National University of Singapore) for providing *pER22-SUC2:FT-GFP ft-10* lines. Y.X. was in part supported by a Chinese Scholarship Council PhD stipend. This article is part of a project (PLAMORF) that has received funding from the European Research Council (ERC) under the European Union’s Horizon 2020 research and innovation program (Grant agreement No. 810131).

## AUTHOR CONTRIBUTIONS

Y.X. with F.K. designed the experiments, Y.X. performed most of the experiments, produced transgenic and grafted plants, and performed RIP experiments, and supported by Y.Z. analyzed the RIP RNAseq data. L.Y. suggested experiments. Y.H. and E.M. performed wild-type grafts and control experiments. A.S. contributed vectors and transgenic plants and performed some initial grafting experiments. S.O. and L.W. performed thermophoretic protein-RNA interaction assays. S.O. identified ALY protein in the B. napus phloem proteome. F.A. performed statistical analysis on mobile transcripts. S.G. analyzed transcriptomes and DEGs. E.S., Y.H., A.S., F.A., and S.G. discussed and commented on the manuscript text. Y.X. and F.K., supported by E.S. and F.A., wrote the manuscript.

## SUPPORTING INFORMATION

## Supporting figures

Figure S1: RT-PCR detection of *YFP* in *35S::YFP*-expressing grafts and CLSM images of YFP- TCTP1 in grafted *A. thaliana* plants.

Figure S2: Fluorescent signal detected in 35S::YFP-TCTP1 (aly4)/aly4 grafts and ALY2/4-GFP transgenic plants.

Figure S3: RIP sample controls showing enrichment of ALY2-GFP and ALY4-GFP proteins in samples used for deep sequencing.

Figure S4: *ALY2* and *ALY4* mRNAs are not mobile in grafted plants.

Figure S5: Endogenous *GRP7* mRNA is mobile in vegetative and flowering grafted *dnmt2 nsun2b* mutants.

Figure S6: Flowering time differences between wild-type (Col-0) and mutant plants.

## Supporting tables

Table S1: Expression changes of selected genes.

## Supporting data

Data S1: Overview of RNAseq samples used in this study.

Data S2: ALY2 and ALY4 RNA IP data.

Data S3: DEGs found in vegetative stage *dnmt2 nsun2b*, *aly2* or *aly4* mutants *and dnmt2 nsun2b* mutants committed to flowering.

Data S4: PCR and cloning primers used in the study.

## REFERENCES

Amara, U., Hu, J., Cai, J. and Kang, H. (2023) FLK is an mRNA m(6)A reader that regulates floral transition by modulating the stability and splicing of FLC in Arabidopsis. Mol Plant, 16, 919–929.

Apelt, F., Breuer, D., Olas, J.J., Annunziata, M.G., Flis, A., Nikoloski, Z., Kragler, F. and Stitt, M. (2017) Circadian, Carbon, and Light Control of Expansion Growth and Leaf Movement. Plant Physiol, 174, 1949–1968.

Arabidopsis Interactome Mapping, C. (2011) Evidence for network evolution in an Arabidopsis interactome map. Science, 333, 601–607.

Bailey, T.L. and Grant, C.E. (2021) SEA: Simple Enrichment Analysis of motifs. bioRxiv, 2021.2008.2023.457422.

Banerjee, A.K., Chatterjee, M., Yu, Y., Suh, S.-G., Miller, W.A. and Hannapel, D.J. (2006) Dynamics of a Mobile RNA of Potato Involved in a Long-Distance Signaling Pathway. The Plant Cell, 18, 3443–3457.

Burgess, A.L., David, R. and Searle, I.R. (2015) Conservation of tRNA and rRNA 5- methylcytosine in the kingdom Plantae. BMC plant biology, 15, 199.

Canto, T., Uhrig, J.F., Swanson, M., Wright, K.M. and MacFarlane, S.A. (2006) Translocation of Tomato bushy stunt virus P19 protein into the nucleus by ALY proteins compromises its silencing suppressor activity. J Virol, 80, 9064–9072.

Cao, S.-K., Du, H., Li, Y., Zhang, J. and Liu, R. (2025) MicroScale thermophoresis (MST): precise analysis of molecular interactions in plants. Trends in Plant Science.

Chen, X., Li, A., Sun, B.-F., Yang, Y., Han, Y.-N., Yuan, X., Chen, R.-X., Wei, W.-S., Liu, Y. and Gao, C.-C.J.N.c.b. (2019) 5-methylcytosine promotes pathogenesis of bladder cancer through stabilizing mRNAs. 21, 978–990.

Cheng, S.L.H., Xu, H., Ng, J.H.T. and Chua, N.-H. (2023) Systemic movement of long non-coding RNA ELENA1 attenuates leaf senescence under nitrogen deficiency. Nature Plants.

Cui, X., Liang, Z., Shen, L., Zhang, Q., Bao, S., Geng, Y., Zhang, B., Leo, V., Vardy, L.A. and Lu, T. (2017) 5-Methylcytosine RNA methylation in Arabidopsis thaliana. Molecular plant, 10, 1387–1399.

David, R., Burgess, A., Parker, B., Li, J., Pulsford, K., Sibbritt, T., Preiss, T. and Searle, I.R. (2017) Transcriptome-Wide Mapping of RNA 5-Methylcytosine in Arabidopsis mRNAs and Noncoding RNAs. Plant Cell, 29, 445–460.

Davis, A.M., Hall, A., Millar, A.J., Darrah, C. and Davis, S.J. (2009) Protocol: Streamlined sub-protocols for floral-dip transformation and selection of transformants in Arabidopsis thaliana. Plant Methods, 5, 3.

Dobin, A., Davis, C.A., Schlesinger, F., Drenkow, J., Zaleski, C., Jha, S., Batut, P., Chaisson, M. and Gingeras, T.R. (2013) STAR: ultrafast universal RNA-seq aligner. Bioinformatics 29, 15–21.

Duan, H.C., Wei, L.H., Zhang, C., Wang, Y., Chen, L., Lu, Z., Chen, P.R., He, C. and Jia, G. (2017) ALKBH10B Is an RNA N(6)-Methyladenosine Demethylase Affecting Arabidopsis Floral Transition. Plant Cell, 29, 2995–3011.

Ehrnsberger, H.F., Grasser, M. and Grasser, K.D. (2019) Nucleocytosolic mRNA transport in plants: export factors and their influence on growth and development. J Exp Bot, 70, 3757–3763.

Fernandes, R., Ostendorp, A., Ostendorp, S., Mehrmann, J., Falke, S., Graewert, M.A., Weingartner, M., Kehr, J. and Hoth, S. (2023) Structural and functional analysis of a plant nucleolar RNA chaperone-like protein. Sci Rep, 13, 9656.

Gao, Y. and Fang, J. (2021) RNA 5-methylcytosine modification and its emerging role as an epitranscriptomic mark. RNA Biol, 18, 117–127.

Ghate, T.H., Sharma, P., Kondhare, K.R., Hannapel, D.J. and Banerjee, A.K. (2017) The mobile RNAs, StBEL11 and StBEL29, suppress growth of tubers in potato. Plant Molecular Biology, 93, 563–578.

Goll, M.G., Kirpekar, F., Maggert, K.A., Yoder, J.A., Hsieh, C.L., Zhang, X., Golic, K.G., Jacobsen, S.E. and Bestor, T.H. (2006) Methylation of tRNAAsp by the DNA methyltransferase homolog Dnmt2. Science, 311, 395–398.

Gutiérrez-Mireles, E.R., Páez-Franco, J.C., Rodríguez-Ruíz, R., Germán-Acacio, J.M., Ló pez-Aquino, M.C. and Gutiérrez-Aguilar, M. (2023) An Arabidopsis mutant line lacking the mitochondrial calcium transport regulator MICU shows an altered metabolite profile. Plant Signaling & Behavior, 18, 2271799.

Haywood, V., Yu, T.S., Huang, N.C. and Lucas, W.J. (2005) Phloem long-distance trafficking of GIBBERELLIC ACID-INSENSITIVE RNA regulates leaf development. Plant J, 42, 49–68.

Hou, N., Li, C., He, J., Liu, Y., Yu, S., Malnoy, M., Mobeen Tahir, M., Xu, L., Ma, F. and Guan, Q. (2022) MdMTA-mediated m(6) A modification enhances drought tolerance by promoting mRNA stability and translation efficiency of genes involved in lignin deposition and oxidative stress. New Phytol, 234, 1294–1314.

Huang, N.C. and Yu, T.S. (2009) The sequences of Arabidopsis GA-INSENSITIVE RNA constitute the motifs that are necessary and sufficient for RNA long-distance trafficking. Plant J, 59, 921–929.

Khan, J.A., Wang, Q., Sjolund, R.D., Schulz, A. and Thompson, G.A. (2007) An early nodulin-like protein accumulates in the sieve element plasma membrane of Arabidopsis. Plant Physiol, 143, 1576–1589.

Kiervel, D., Boissinot, S., Piccini, C., Scheidecker, D., Villeroy, C., Gilmer, D., Brault, V. and Ziegler-Graff, V. (2024) Arabidopsis RNA–binding proteins interact with viral structural proteins and modify turnip yellows virus accumulation. Plant Physiology, 197.

Kitagawa, M., Wu, P., Balkunde, R., Cunniff, P. and Jackson, D. (2022) An RNA exosome subunit mediates cell-to-cell trafficking of a homeobox mRNA via plasmodesmata. 375, 177–182.

Koroleva, O.A., Calder, G., Pendle, A.F., Kim, S.H., Lewandowska, D., Simpson, C.G., Jones, I.M., Brown, J.W. and Shaw, P.J. (2009) Dynamic behavior of Arabidopsis eIF4A-III, putative core protein of exon junction complex: fast relocation to nucleolus and splicing speckles under hypoxia. Plant Cell, 21, 1592–1606.

Kubina, J., Geldreich, A., Gales, J.P., Baumberger, N., Bouton, C., Ryabova, L.A., Grasser, K.D., Keller, M. and Dimitrova, M. (2021) Nuclear export of plant pararetrovirus mRNAs involves the TREX complex, two viral proteins and the highly structured 5’ leader region. Nucleic Acids Res, 49, 8900–8922.

Li, S., Wang, X., Xu, W., Liu, T., Cai, C., Chen, L., Clark, C.B. and Ma, J. (2021) Unidirectional movement of small RNAs from shoots to roots in interspecific heterografts. Nature Plants, 7, 50–59.

Li, X., Wang, C., Chen, Y., Liu, W., Zhang, M., Wang, N., Xiang, C., Gao, L., Dong, Y. and Zhang, W. (2024) m5C and m6A modifications regulate the mobility of pumpkin CHOLINE KINASE 1 mRNA under chilling stress. Plant Physiology, 197.

Liao, Y., Smyth, G.K. and Shi, W. (2014) featureCounts: an efficient general purpose program for assigning sequence reads to genomic features. Bioinformatics, 30, 923–930.

Liu, L., Li, C., Teo, Z.W.N., Zhang, B. and Yu, H. (2019a) The MCTP-SNARE Complex Regulates Florigen Transport in Arabidopsis. Plant Cell, 31, 2475–2490.

Liu, Y., Ma, Y., Salsman, E., Manthey, F.A., Elias, E.M., Li, X. and Yan, C. (2019b) An enrichment method for mapping ambiguous reads to the reference genome for NGS analysis. J Bioinform Comput Biol, 17, 1940012.

Love, M.I., Huber, W. and Anders, S. (2014) Moderated estimation of fold change and dispersion for RNA-seq data with DESeq2. Genome Biology, 15, 550.

Lu, K.-J., Huang, N.-C., Liu, Y.-S., Lu, C.-A. and Yu, T.-S. (2012) Long-distance movement of Arabidopsis FLOWERING LOCUS T RNA participates in systemic floral regulation. RNA biology, 9, 653–662.

Mandel, M.A. and Yanofsky, M.F. (1995) A gene triggering flower formation in Arabidopsis. Nature, 377, 522–524.

Meyer, K.D. and Jaffrey, S.R. (2014) The dynamic epitranscriptome: N6-methyladenosine and gene expression control. Nature Reviews Molecular Cell Biology, 15, 313–326.

Ostendorp, A., Ostendorp, S., Zhou, Y., Chaudron, Z., Wolffram, L., Rombi, K., von Pein, L., Falke, S., Jeffries, C.M., Svergun, D.I., Betzel, C., Morris, R.J., Kragler, F. and Kehr, J. (2022) Intrinsically disordered plant protein PARCL colocalizes with RNA in phase-separated condensates whose formation can be regulated by mutating the PLD. J Biol Chem, 298, 102631.

Ostendorp, A., Pahlow, S., Krüßel, L., Hanhart, P., Garbe, M.Y., Deke, J., Giavalisco, P. and Kehr, J. (2017) Functional analysis of Brassica napus phloem protein and ribonucleoprotein complexes. New Phytol, 214, 1188–1197.

Paajanen, P., Tomkins, M., Hoerbst, F., Veevers, R., Heeney, M., Thomas, H.R., Apelt, F., Saplaoura, E., Gupta, S., Frank, M., Walther, D., Faulkner, C., Kehr, J., Kragler, F. and Morris, R.J. (2025) Re-analysis of mobile mRNA datasets raises questions about the extent of long-distance mRNA communication. Nature Plants.

Park, N.I., Yeung, E.C. and Muench, D.G. (2009) Mago Nashi is involved in meristem organization, pollen formation, and seed development in Arabidopsis. Plant Sci, 176, 461–469.

Paultre, D.S.G., Gustin, M.P., Molnar, A. and Oparka, K.J. (2016) Lost in Transit: Long-Distance Trafficking and Phloem Unloading of Protein Signals in Arabidopsis Homografts. Plant Cell, 28, 2016–2025.

Pfaff, C., Ehrnsberger, H.F., Flores-Tornero, M., Sorensen, B.B., Schubert, T., Langst, G., Griesenbeck, J., Sprunck, S., Grasser, M. and Grasser, K.D. (2018a) ALY RNA-Binding Proteins Are Required for Nucleocytosolic mRNA Transport and Modulate Plant Growth and Development. Plant Physiol, 177, 226–240.

Pfaff, C., Ehrnsberger, H.F., Flores-Tornero, M., Sørensen, B.B., Schubert, T., Längst, G., Griesenbeck, J., Sprunck, S., Grasser, M. and Grasser, K.D. (2018b) ALY RNA-binding proteins are required for nucleocytosolic mRNA transport and modulate plant growth and development. Plant physiology, 177, 226–240.

Prall, W., Sheikh, A.H., Bazin, J., Bigeard, J., Almeida-Trapp, M., Crespi, M., Hirt, H. and Gregory, B.D. (2023) Pathogen-induced m6A dynamics affect plant immunity. The Plant Cell.

Rappsilber, J., Mann, M. and Ishihama, Y. (2007) Protocol for micro-purification, enrichment, pre-fractionation and storage of peptides for proteomics using StageTips. Nat Protoc, 2, 1896–1906.

Ross-Elliott, T.J., Jensen, K.H., Haaning, K.S., Wager, B.M., Knoblauch, J., Howell, A.H., Mullendore, D.L., Monteith, A.G., Paultre, D., Yan, D., Otero, S., Bourdon, M., Sager, R., Lee, J.Y., Helariutta, Y., Knoblauch, M. and Oparka, K.J. (2017) Phloem unloading in Arabidopsis roots is convective and regulated by the phloem-pole pericycle. Elife, 6.

Schladt, T.M., Miras, M., Ejike, J.O., Pottier, M., Xi, L., Restrepo-Escobar, A., Nakamura, M., Pütz, N., Hänsch, S., Gao, C., Engelhorn, J., Dickmanns, M., Davis, G.V., Dalal, A., Gombos, S., Lange, R., Simon, R., Schulze, W.X. and Frommer, W.B. (2025) Identification of nuclear pore proteins at plasmodesmata: potential role in intercellular transport?: eLife Sciences Publications, Ltd.

Scutenaire, J., Deragon, J.M., Jean, V., Benhamed, M., Raynaud, C., Favory, J.J., Merret, R. and Bousquet-Antonelli, C. (2018) The YTH Domain Protein ECT2 Is an m(6)A Reader Required for Normal Trichome Branching in Arabidopsis. Plant Cell, 30, 986–1005.

Seidel, S.A., Dijkman, P.M., Lea, W.A., van den Bogaart, G., Jerabek-Willemsen, M., Lazic, A., Joseph, J.S., Srinivasan, P., Baaske, P., Simeonov, A., Katritch, I., Melo, F.A., Ladbury, J.E., Schreiber, G., Watts, A., Braun, D. and Duhr, S. (2013) Microscale thermophoresis quantifies biomolecular interactions under previously challenging conditions. Methods, 59, 301–315.

Shen, L., Liang, Z., Gu, X., Chen, Y., Teo, Z.W., Hou, X., Cai, W.M., Dedon, P.C., Liu, L. and Yu, H. (2016) N(6)-Methyladenosine RNA Modification Regulates Shoot Stem Cell Fate in Arabidopsis. Dev Cell, 38, 186–200.

Simon, R., Igeño, M.I. and Coupland, G. (1996) Activation of floral meristem identity genes in Arabidopsis. Nature, 384, 59–62.

Song, P., Yang, J., Wang, C., Lu, Q., Shi, L., Tayier, S. and Jia, G. (2021) Arabidopsis N6- methyladenosine reader CPSF30-L recognizes FUE signals to control polyadenylation site choice in liquid-like nuclear bodies. Molecular Plant, 14, 571–587.

Strauch, C.J., Sprotte, N., Peña Lozano, E., Boutant, E., Amari, K., Ostendorp, S., Ostendorp, A., Kehr, J. and Niehl, A. (2025) Studies on the Japanese soil-borne wheat mosaic virus movement protein highlight its ability to bind plant RNA. Virology Journal, 22, 134.

Tang, Y., Gao, C.C., Gao, Y., Yang, Y., Shi, B., Yu, J.L., Lyu, C., Sun, B.F., Wang, H.L., Xu, Y., Yang, Y.G. and Chong, K. (2020) OsNSUN2-Mediated 5-Methylcytosine mRNA Modification Enhances Rice Adaptation to High Temperature. Dev Cell, 53, 272–286.e277.

Thieme, C.J., Rojas-Triana, M., Stecyk, E., Schudoma, C., Zhang, W., Yang, L., Mi√± ambres, M., Walther, D., Schulze, W.X., Paz-Ares, J., Scheible, W.-R.d. and Kragler, F. (2015) Endogenous Arabidopsis messenger RNAs transported to distant tissues. Nature Plants, 1, 15025.

Uhrig, J.F., Canto, T., Marshall, D. and MacFarlane, S.A. (2004) Relocalization of nuclear ALY proteins to the cytoplasm by the tomato bushy stunt virus P19 pathogenicity protein. Plant Physiol, 135, 2411–2423.

Wang, M. and Teng, Y. (2018) Genome-wide identification and analysis of MICU genes in land plants and their potential role in calcium stress. Gene, 670, 174–181.

Wei, L.H., Song, P., Wang, Y., Lu, Z., Tang, Q., Yu, Q., Xiao, Y., Zhang, X., Duan, H.C. and Jia, G. (2018) The m(6)A Reader ECT2 Controls Trichome Morphology by Affecting mRNA Stability in Arabidopsis. Plant Cell, 30, 968–985.

Wickramasinghe, V.O. and Laskey, R.A. (2015) Control of mammalian gene expression by selective mRNA export. Nature Reviews Molecular Cell Biology, 16, 431–442.

Wu, T., Hu, E., Xu, S., Chen, M., Guo, P., Dai, Z., Feng, T., Zhou, L., Tang, W., Zhan, L., Fu, X., Liu, S., Bo, X. and Yu, G. (2021) clusterProfiler 4.0: A universal enrichment tool for interpreting omics data. Innovation (Camb), 2, 100141.

Xia, C., Zheng, Y., Huang, J., Zhou, X., Li, R., Zha, M., Wang, S., Huang, Z., Lan, H., Turgeon, R., Fei, Z. and Zhang, C. (2018) Elucidation of the Mechanisms of Long-Distance mRNA Movement in a Nicotiana benthamiana/Tomato Heterograft System. Plant Physiology, 177, 745–758.

Xue, C., Zhao, Y. and Li, L. (2020) Advances in RNA cytosine-5 methylation: detection, regulatory mechanisms, biological functions and links to cancer. Biomarker Research, 8, 43.

Yang, L., Machin, F., Wang, S., Saplaoura, E. and Kragler, F. (2023a) Heritable transgene-free genome editing in plants by grafting of wild-type shoots to transgenic donor rootstocks. Nat Biotechnol, 41, 958–967.

Yang, L., Perrera, V., Saplaoura, E., Apelt, F., Bahin, M., Kramdi, A., Olas, J., Mueller-Roeber, B., Sokolowska, E., Zhang, W., Li, R., Pitzalis, N., Heinlein, M., Zhang, S., Genovesio, A., Colot, V. and Kragler, F. (2019) m(5)C Methylation Guides Systemic Transport of Messenger RNA over Graft Junctions in Plants. Curr Biol, 29, 2465–2476 e2465.

Yang, L., Zhou, Y., Wang, S., Xu, Y., Ostendorp, S., Tomkins, M., Kehr, J., Morris, R.J. and Kragler, F. (2023b) Noncell-autonomous HSC70.1 chaperone displays homeostatic feedback regulation by binding its own mRNA. New Phytol, 237, 2404–2421.

Yang, X., Yang, Y., Sun, B.-F., Chen, Y.-S., Xu, J.-W., Lai, W.-Y., Li, A., Wang, X., Bhattarai, D.P. and Xiao, W. (2017) 5-methylcytosine promotes mRNA export—NSUN2 as the methyltransferase and ALYREF as an m5C reader. Cell Research, 27, 606–625.

Yang, Y., Mao, L., Jittayasothorn, Y., Kang, Y., Jiao, C., Fei, Z. and Zhong, G.-Y. (2015) Messenger RNA exchange between scions and rootstocks in grafted grapevines. BMC plant biology, 15, 251.

Zhang, F., Zhang, Y.C., Liao, J.Y., Yu, Y., Zhou, Y.F., Feng, Y.Z., Yang, Y.W., Lei, M.Q., Bai, M., Wu, H. and Chen, Y.Q. (2019) The subunit of RNA N6-methyladenosine methyltransferase OsFIP regulates early degeneration of microspores in rice. PLoS Genet, 15, e1008120.

Zhang, W., Kollwig, G., Stecyk, E., Apelt, F., Dirks, R. and Kragler, F. (2014) Graft-transmissible movement of inverted-repeat-induced siRNA signals into flowers. Plant J, 80, 106–121.

Zhang, W., Thieme, C.J., Kollwig, G., Apelt, F., Yang, L., Winter, N., Andresen, N., Walther, D. and Kragler, F. (2016) tRNA-Related Sequences Trigger Systemic mRNA Transport in Plants. Plant Cell, 28, 1237–1249.

Zhou, L., Tang, R., Li, X., Tian, S., Li, B. and Qin, G. (2021) N6-methyladenosine RNA modification regulates strawberry fruit ripening in an ABA-dependent manner. Genome Biology, 22, 168.

Zhou, Z., Luo, M.J., Straesser, K., Katahira, J., Hurt, E. and Reed, R. (2000) The protein Aly links pre-messenger-RNA splicing to nuclear export in metazoans. Nature, 407, 401–405.

