## Supplemental Figure S1 for "Developmental Control of Long-Distance mRNA Transport Involves m^5^C RNA Methylation and ALYREF Nuclear Export Factors"

### Xu et al. 2026 Figure S1

**A** RT-PCR assays showing that *YFP* mRNA is not graft-mobile

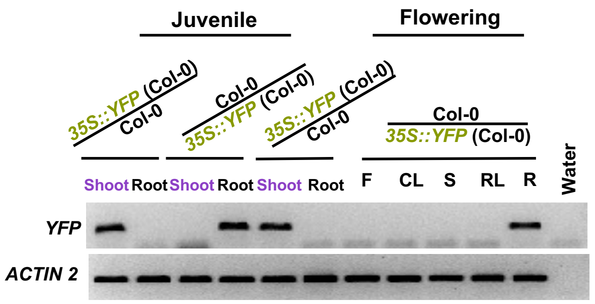


**B** CLSM images and YFP-TCTP1 presence in phloem pericycle cells

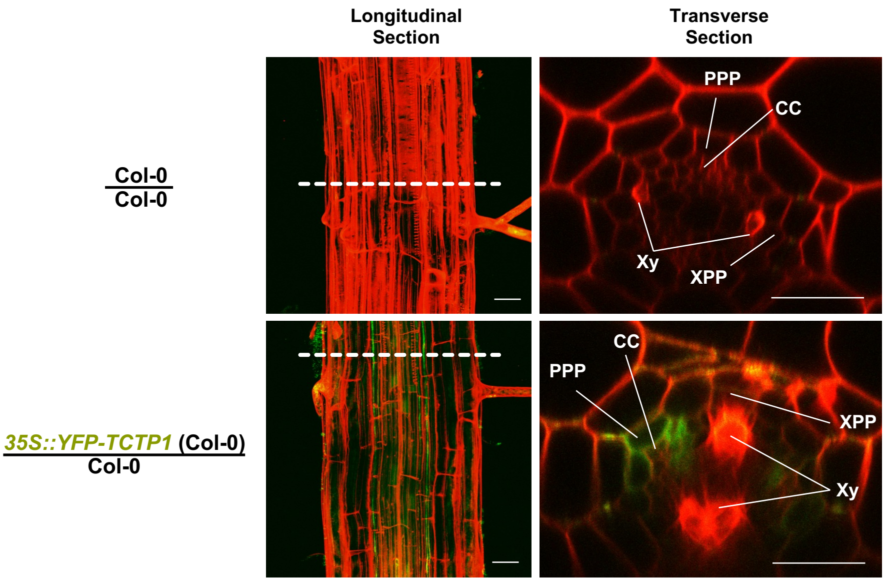
