## Supplementary figures and images for "Developmental Control of Long-Distance mRNA Transport Involves m^5^C RNA Methylation and ALYREF Nuclear Export Factors"

### Supplemental Figure S2

#### Xu et al. 2026 Figure S2

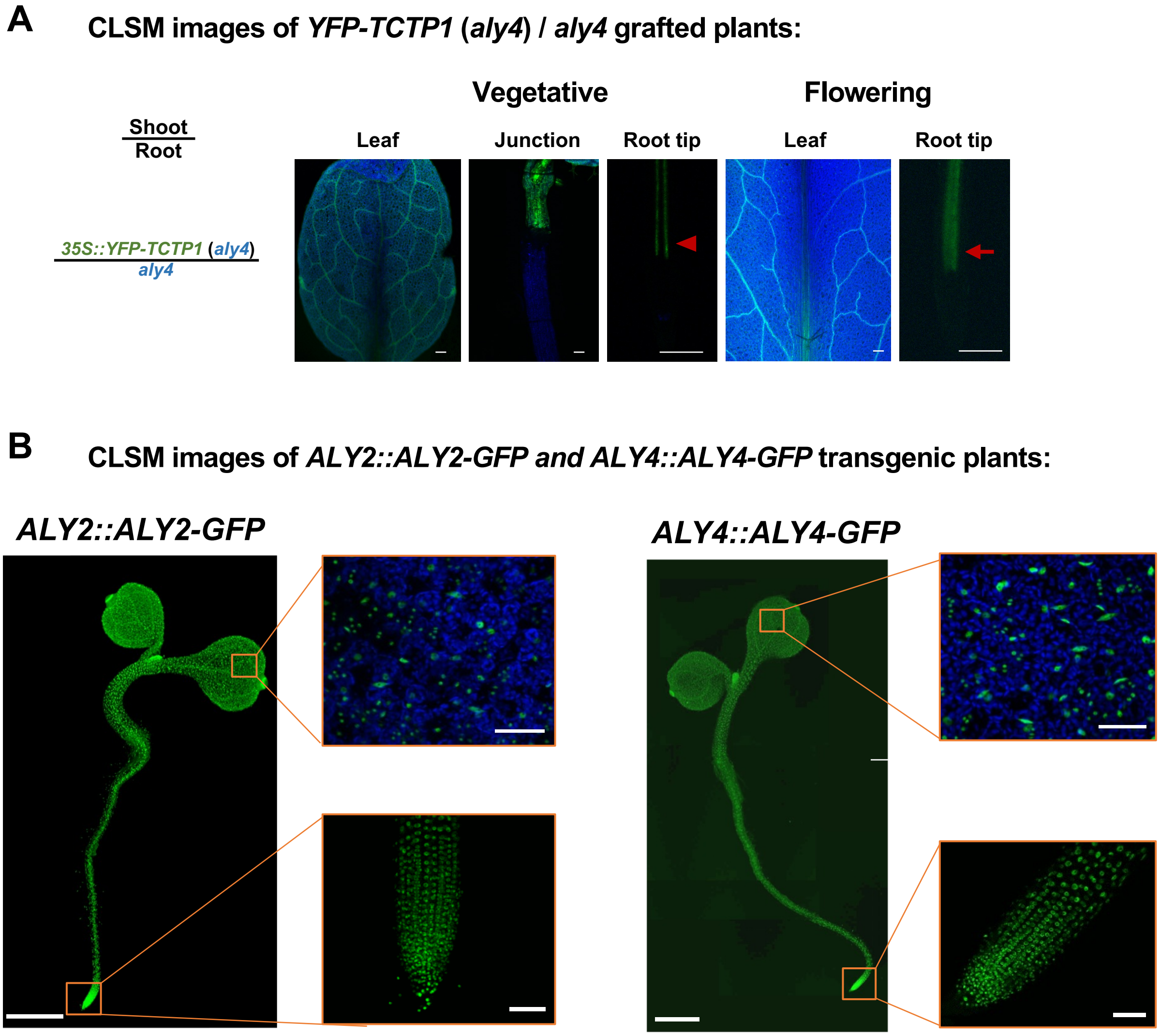

### Supplemental Figure S3

#### Xu et al. 2026 Figure S3

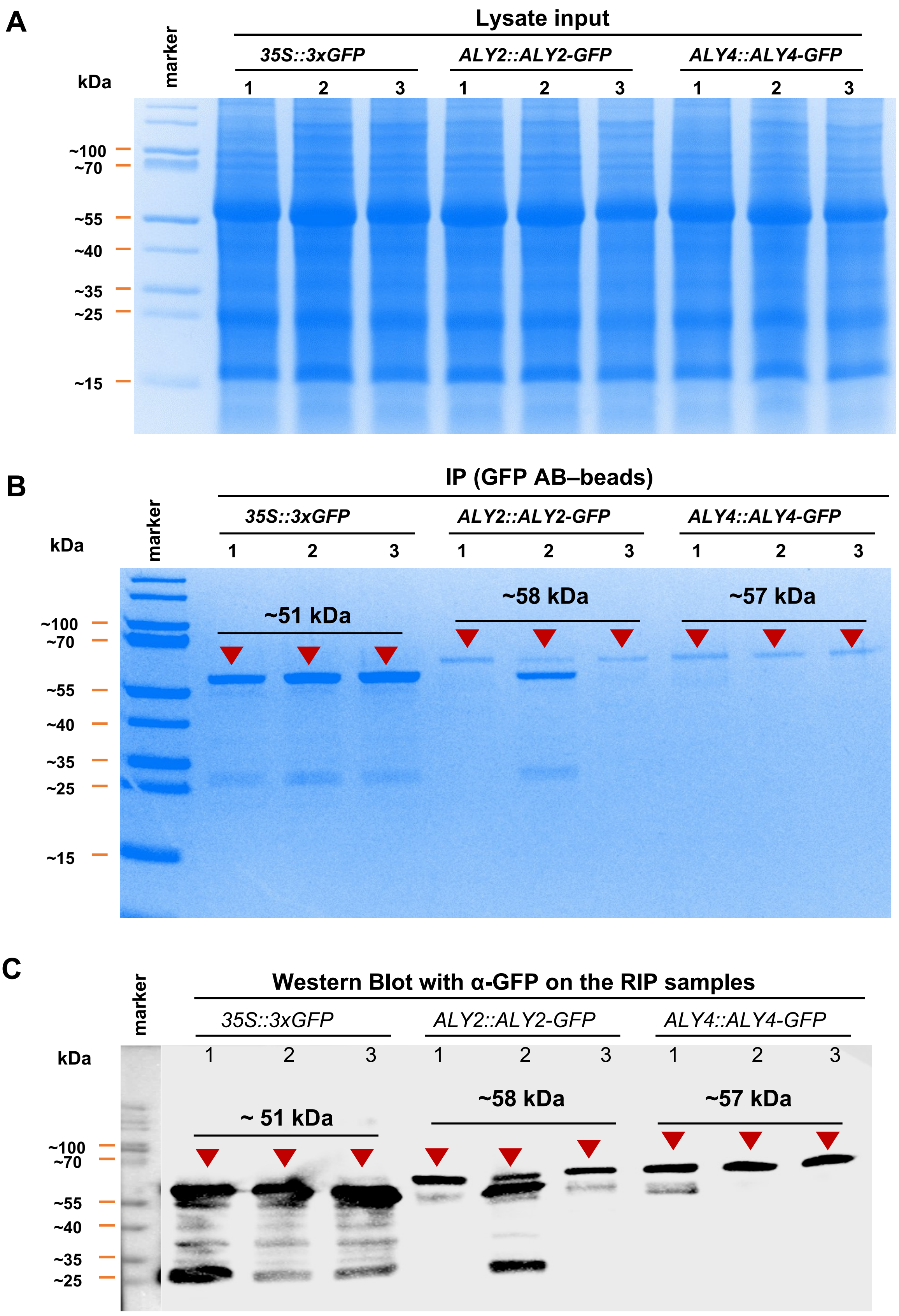

### Supplemental Figure S4

#### Xu et al. 2026 Figure S4

##### RT-PCR assays to confirm lack of *ALY2 and ALY4* transcript mobility:

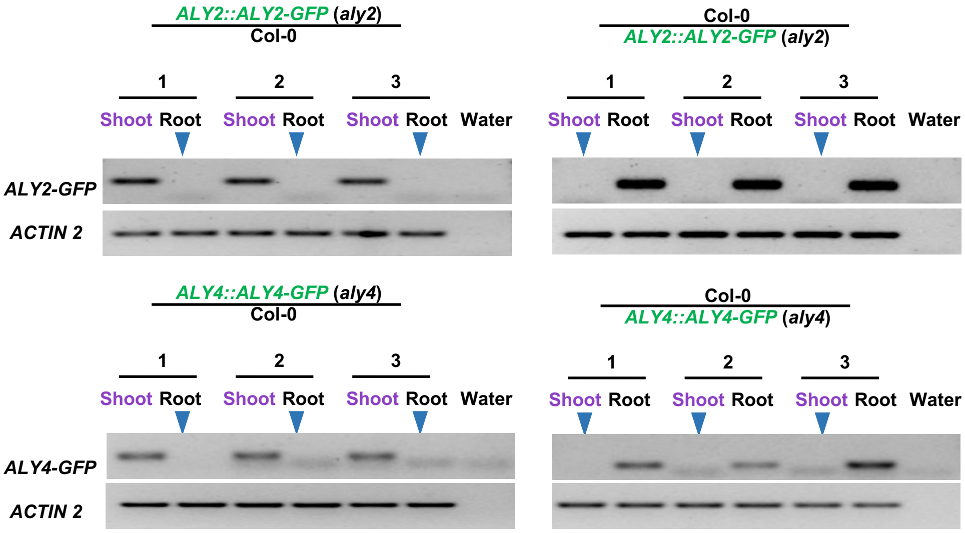
