## Supplemental Figure S5 for "Developmental Control of Long-Distance mRNA Transport Involves m^5^C RNA Methylation and ALYREF Nuclear Export Factors"

### Xu et al. 2026 Figure S5

#### RT-PCR assays indicate *GRP7* transcript mobility in vegetative and flowering *dnmt2 nsun2b* mutants

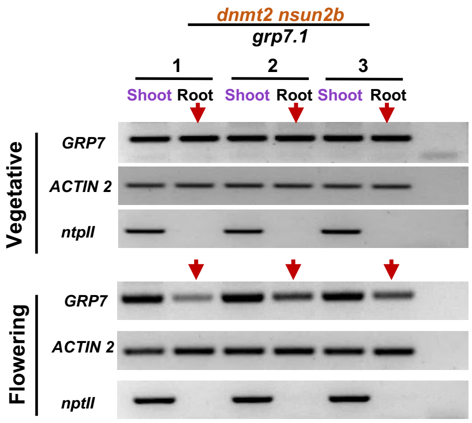
