## Supplemental Figure S6 for "Developmental Control of Long-Distance mRNA Transport Involves m^5^C RNA Methylation and ALYREF Nuclear Export Factors"

### Xu et al. 2026 Figure S6

**Flowering time (rosette leaf number & bolding time) analysis of *mutants vs.* wild-type:**

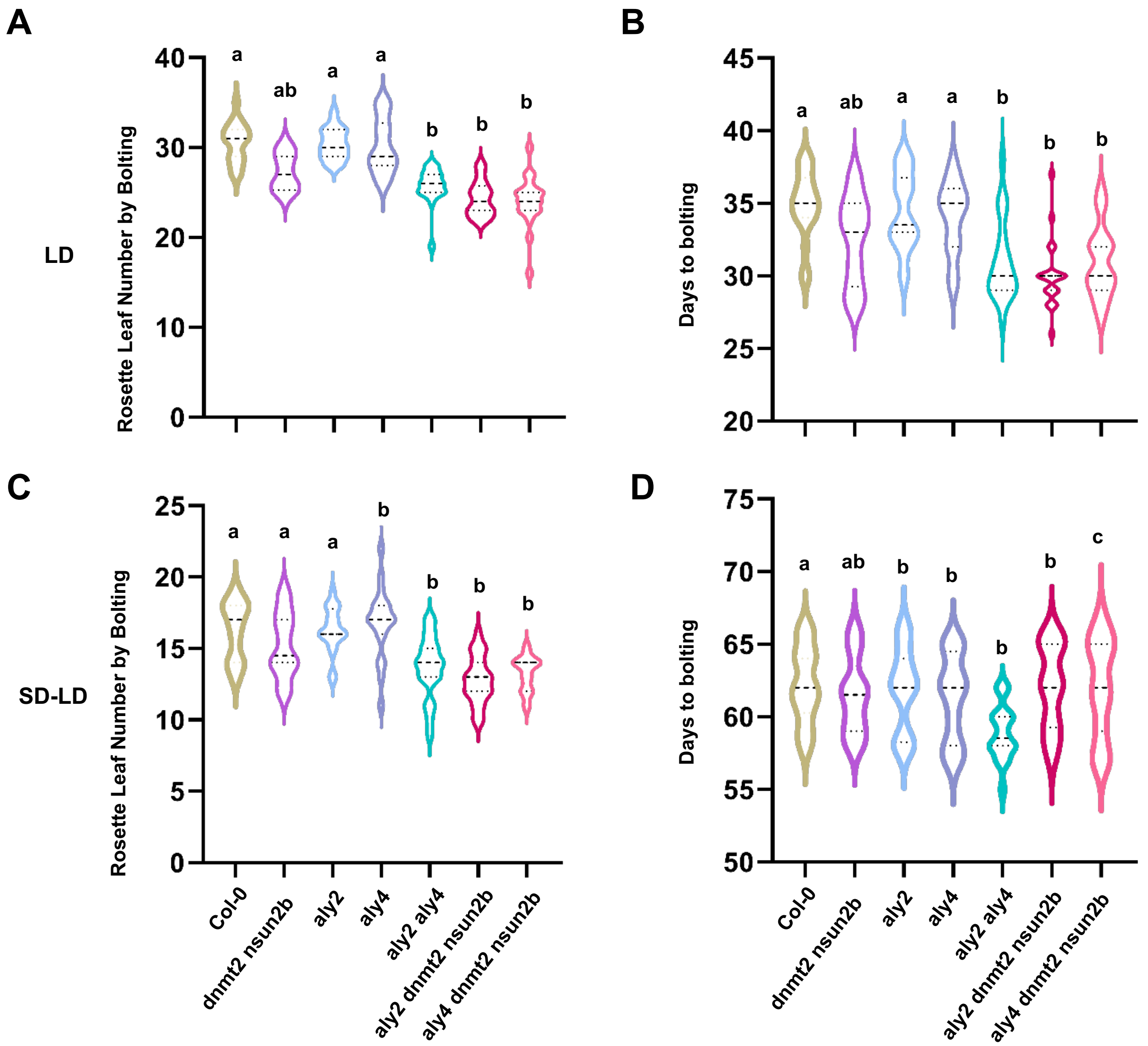
