## Supplemental Data S4 for "Developmental Control of Long-Distance mRNA Transport Involves m^5^C RNA Methylation and ALYREF Nuclear Export Factors"

### Xu et al. 2026

### Data S4

### Table of oligonucleotide sequences used in the study

| **Name of primer** | **Primer sequences** | **Size of PCR fragment** |
| --- | --- | --- |
| *ACTIN2*  (AT3G18780) | FK424-F 5’-GGAAGGATCTGTACGGTAAC-3’ | 245 bp |
|  | FK425-R 5’-TGTGAACGATTCCTGGACCT-3’ |  |
| *YFP-TCTP1* | FK1210-F 5’-GCTTCGAATTCTGCAGTCGAC-3’ | 137 bp |
|  | FK1211-R 5’-GGGAAAGAGTCAGACAGAAGCTC-3’ |  |
| *YFP* | FK1166-F 5’-CAGCACGACTTCTTCAAGTCC-3’ | 274 bp |
|  | FK1167-R 5’-TGTTGTGGCGGATCTTGAAGT-3’ |  |
| *HSC70.1-GFP* | FK1328-F 5’-GCTGGTGGTGAAGCCGGTG-3’ | 280 bp |
|  | FK1329-R 5’-CGCTGAACTTGTGGCCGTTT-3’ |  |
| *AP1*  (AT1G69120) | FK1688-F 5’-CAGCACCAAATCCAGCATCCTTAC-3’ | 154 bp |
|  | FK1689-R 5’-AGCAGCCAAGGTTGCAGTTGTAAA-3’ |  |
| *GRP7*  (AT2G21660) | FK1373-F 5’-ATGGCGTCCGGTGATGTTGA-3’ | 271 bp |
|  | FK1374-R 5’-CACCGCTTCCTCGTGACTGA-3’ |  |
| *BAR* | FK966-F 5’-CAGGAACCGCAGGAGTGGA-3’ | 407 bp |
|  | FK967-R 5’-CCAGAAACCCACGTCATGCC-3’ |  |
| *Hyg* | FK1012-F 5’-ATGAAAAAGCCTGAACTCACC-3’ | 114 bp |
|  | FK1013-R 5’-GCTGAAAGCACGAGATTCTTC-3’ |  |
| *NPTII* | FK1339-F 5’-AGAGGCTATTCGGCTATGACTGG-3’ | 450 bp |
|  | FK1340-R 5’-ATCGCCATGGGTCACGACGAGAT-3’ |  |
| *GRP7-YFP* | FK1907-F 5’-AGCTACGGTGGTGGAAGACGTG-3’ | 208 bp |
|  | FK1912-R 5’-ACTTGTGGCCGTTTACGTCGCC-3’ |  |
| *T7_MICU* | F 5’-TAATACGACTCACTATAGGGGGGCAAATCTAAAAGATAAGGTTC-3’ |  |
|  | R 5’-CTTAAGATTATATACAAAGATATGTATC-3’ |  |
| *T7_GRP7* | F 5’-TAATACGACTCACTATAGGGGCTTCGTCTACATCGTTCTACAC-3’ |  |
|  | R 5’-AGATGATAGAATCAATCAACAGAG-3’ |  |
| *T7_HSC70.1* | F 5’-TAATACGACTCACTATAGGGGACTGAATAATGCCAACGTGTAC-3’ |  |
|  | R 5’-AAGTAATTTTTATGTTATGCCATTTGGG-3’ |  |
| *T7_TCTP1* | F 5’-TAATACGACTCACTATAGGGGACTGAATAATGCCAACGTGTAC-3’ |  |
|  | R 5’-TCAGCACTTGACCTCCTTCAAACCATG-3’ |  |
| *GRP7_Promoter*  (SalI-BamHI) | F 5'-AAAAGTCGACAAATCTTCTCCTTCATCACCGCA-3' | 2253 bp |
|  | R 5'-AAAAGGATCCTGAAATTTGAAAAGAAGATCTAAGGG-3' |  |
| *GRP7_CDS_linker_ (-Stop)* (BamHI-StuI) | F 5'-GGGGGGATCCATGGCGTCCGGTGATGTTGAGTATCGG-3' | 578 bp |
|  | R 5'-GGGGAGGCCTCGCAGCAGCCGCAGCAGCAGCCGCAGCAGCCCA  TCCTCCACCACCACCGCTTCC-3' |  |
| *YFP*  (StuI-EagI) | F 5’-GGGGAGGCCTATGGTGAGCAAGGGCGAGGAGCTGTTC-3’ | 740 bp |
|  | R 5’-GGGGCGGCCGTTACTTGTACAGCTCGTCCATGCCGAGAG-3’ |  |
| *GRP7_3’UTR*  (EagI-XhoI) | F 5'-CCCCCGGCCGTTCCTTTAATTAGGTTTGGGATTACC-3' | 496 bp |
|  | R 5'-AAAACTCGAGTGTAACATGCATAAGTTGAGTTGG-3' |  |
| *Oligo dT* | FK532 5’-TTTTTTTTTTTTTTTTTTTTVN-3’ |  |
