## Supplemental Table S1 for "Developmental Control of Long-Distance mRNA Transport Involves m^5^C RNA Methylation and ALYREF Nuclear Export Factors"

**Table S1**. Significant expression changes of the *TCTP1*, *GRP7*, and *HSC70.1* mobile transcripts and of *ALY1, ALY3, and* *TRM4A* in the grafted *nsun2b dnmt2* and *aly4* and *aly2* mutants compared to grafted wild-type (Col-0) (log2FC>+/-0.5, padj<0.05; Overall data see: *SI Appendix* Data S4).

|  | **Vegetative** | | | | | **Flowering** |
| --- | --- | --- | --- | --- | --- | --- |
| Gene | Col -0 vs. *aly2* (shoot) | Col -0 vs. *aly4* (shoot)* | Col-0 vs. *nsun2b dnmt2* (shoot)* | *nsun2b dnmt2* vs. *aly2* (shoot) | *nsun2b dnmt2* vs. *aly4* (shoot)* | Col-0 vs.*dnmt2 nsun2b* (shoot) |
| *ALY1 (*AT5G59950) | no | no | no | no | no | no |
| *ALY2 (*AT5G02530) | n/a | no | no | n/a | no | no |
| *ALY3 (*AT1G66260) | no | no | no | no | no | no |
| *ALY4*  *(*At5G37720) | no | n/a | no | no | n/a | no |
| *DNMT2 (*AT5G25480) | no | no | n/a | n/a | n/a | n/a |
| *NSUN2B (*AT2G22400) | no | no | n/a | n/a | n/a | n/a |
| *TRM4A (*AT4G40000) | no | no | no | no | no | no |
| *HSC70.1*  (AT5G02500) | up | no | no | no | no | no |
| *TCTP1*  (AT3G16640) | up | no | no | no | no | no |
| *GRP7*  (AT2G21660) | no | no | no | no | no | no |

* Note that no significant expression changes of mobile *GRP7*, ALY family members, and of *TRM4A* was observed in the *dnmt2 nsun2b* and *aly4* mutants; n/a: not applicable.
